# Myasthenia gravis-specific aberrant neuromuscular gene expression by medullary thymic epithelial cells in thymoma

**DOI:** 10.1101/2021.12.19.473411

**Authors:** Yoshiaki Yasumizu, Naganari Ohkura, Hisashi Murata, Makoto Kinoshita, Soichiro Funaki, Satoshi Nojima, Kansuke Kido, Masaharu Kohara, Daisuke Motooka, Daisuke Okuzaki, Shuji Suganami, Eriko Takeuchi, Yamami Nakamura, Yusuke Takeshima, Masaya Arai, Satoru Tada, Meinoshin Okumura, Eiichi Morii, Yasushi Shintani, Shimon Sakaguchi, Tatsusada Okuno, Hideki Mochizuki

**Affiliations:** Department of Neurology, Graduate School of Medicine, Osaka University, Suita, Osaka, Japan; Department of Experimental Immunology, Immunology Frontier Research Center, Osaka University, Suita, Osaka, Japan; Integrated Frontier Research for Medical Science Division, Institute for Open and Transdisciplinary Research Initiatives (OTRI), Osaka University, Suita, Osaka, Japan; Department of Pathology, Graduate School of Medicine, Osaka University, Suita, Osaka, Japan; Department of General Thoracic Surgery, Graduate School of Medicine, Osaka University, Suita, Osaka, Japan; Genome Information Research Center, Research Institute for Microbial Diseases, Osaka University, Suita, Osaka, Japan; Department of General Thoracic Surgery, National Hospital Organization Toneyama Hospital, Osaka, Japan

**Author notes:** Corresponding authors, Correspondence to Naganari Ohkura and Tatsusada Okuno;.

## Abstract

Myasthenia gravis (MG) is a neurological disease caused by autoantibodies against neuromuscular-associated proteins. While MG is frequently developed in thymoma patients, the etiologic factors for MG are not well understood. Here, by constructing a comprehensive atlas of thymoma using bulk and single-cell RNA-seq, we identified ectopic expression of neuromuscular molecules in MG-associated thymoma (MG-thymoma). These molecules were originated from a distinct subpopulation of medullary thymic epithelial cells (mTECs), which we named neuromuscular mTECs (nmTECs). MG-thymoma also exhibited microenvironments dedicated to autoantibody production, including ectopic germinal center formation, T follicular helper cell accumulation, and type 2 conventional dendritic cell migration. Cell-cell interaction analysis also predicted the interaction between nmTECs and T/B cells via *CXCL12*-*CXCR4*. The enrichment of nmTECs presenting neuromuscular molecules within MG-thymoma was further confirmed by immunohistochemically and by cellular composition estimation from MG-thymoma transcriptome. Altogether, this study suggests that nmTECs play a significant role in MG pathogenesis via ectopic expression of neuromuscular molecules.

## Main

Myasthenia gravis (MG) is the most common disorder of neuromuscular transmission caused by autoantibodies against the motor endplate, such as anti-acetylcholine receptor (AChR) antibodies. MG is often accompanied by thymoma, and thymoma-associated MG (TAMG) is more difficult to manage than other forms of MG because of its frequent crisis, the need for surgery, the difficulty of perioperative management and the need for intense immunotherapies^1^. As the epidemiology, 21% of MG patients experienced thymoma^2^, and 25% of thymoma patients experienced MG^3^, indicating that MG and abnormalities in the thymus are closely related to each other. This is also exemplified by that thymectomy is a well-established treatment for TAMG in addition to immunosuppressive treatments^4^.

Abnormalities of the thymus, in which immature thymocytes differentiate into matured CD4^+^ or CD8^+^ T cells, are frequently associated with a variety of autoimmune diseases, such as pure red cell aplasia and Good syndrome^5^. In the thymus, T cell maturation and selection are conducted by the interaction with antigen-presenting cells, including thymic epithelial cells (TECs), myeloid cells, and B cells^6,7^. The positive selection of functional T cells is mediated by cortical TECs (cTECs), while the negative selection of autoreactive T cells is mediated by medullary TECs (mTECs) presenting self-antigens on MHCs. In line with the critical role of mTECs, a loss of function mutation of *AIRE,* which is an essential transcription factor for producing self-antigens in mTECs^8^, causes systemic autoimmunity called Autoimmune Polyglandular Syndrome Type 1 (APS-1)^9^. In addition, dysregulation of the thymus, including thymoma and thymic hyperplasia, are frequently associated not only with MG but also neurological disorders, including encephalitis, which are caused by a wide range of autoantibodies^10,11,12,13,14,15^ Thus, abnormalities of the thymus are closely associated with the generation of self-reactive autoantibodies, which result in the development of autoimmune diseases.

Given that MG is caused by self-reactive autoantibodies, MG-specific changes within thymoma may be a clue for understanding the pathogenesis of MG. It has been reported so far that the accumulation of neurofilaments, which is expressed in neurons under the normal condition, are highly detected in MG-thymoma^16^. In addition, germinal centers (GCs) and T follicular helper (Tfh) cells, both of which play critical roles in antibody production, are also enriched in MG-thymoma^17,18^. Despite the possible contribution of these changes in MG pathology, the complete picture of MG pathogenesis from abnormalities in thymoma to auto-reactive B cell maturation is still poorly understood due to intra- and inter-individual heterogeneity of the thymus. Therefore, to reveal the complex pathogenicity of MG in thymoma, we integratively analyzed bulk and single-cell transcriptomes of MG-thymoma and found that a distinct subpopulation of mTECs ectopically expressed neuromuscular-associated molecules and would contribute to the pathogenesis of MG via presenting neuromuscular molecules to self-reactive immune cells developed in thymoma.

## Results

### Ectopic expression of neuron-related molecules in MG-thymoma

To characterize MG-specific changes in thymoma comprehensively, we first investigated gene expression profiles from surgically dissected thymoma samples enrolled by The Cancer Genome Atlas (TCGA)^19^ (Fig. 1a). Of the 116 thymoma samples with RNA-seq data, 34 were complicated with MG. In the WHO classifications, which are commonly used for the thymoma staging based on histology, there are six classifications: Type A, AB, B1-B3, and C (i.e., A, spindle cells; AB, mixed spindle cells and lymphocytes; B1, lymphocytes > epithelial cells; B2, mixed lymphocytes and epithelial cells; B3, predominant epithelial cells; and C, carcinoma). In the present data set, MG was associated with multiple types except for type C ( Supplementary Fig. 2a), with its peak at type B2, as previously reported^20^. When we investigated differentially expressed genes between thymoma with and without MG (Supplementary Table 1, Extended Fig. 2b), 93 and 91 genes were identified as upregulated and downregulated genes in MG, respectively. The upregulated genes contained neuromuscular-related molecules; *NEFM*, *RYR3*, *GABRA5*, and immunoreceptors; *PLXNB3*, *IL13RA*. We also observed a slight increase in the acetylcholine receptor *CHRNA1,* which is the main target of autoantibodies in TAMG (*log*_*2*_ *fold change* = 1.07, *P*_*adj*_ = 0.87 with DESeq2^21^; *P* = 0.0051 with two-sided Mann-Whitney *U* test, Supplementary Fig. 2c).

**Figure 1.**
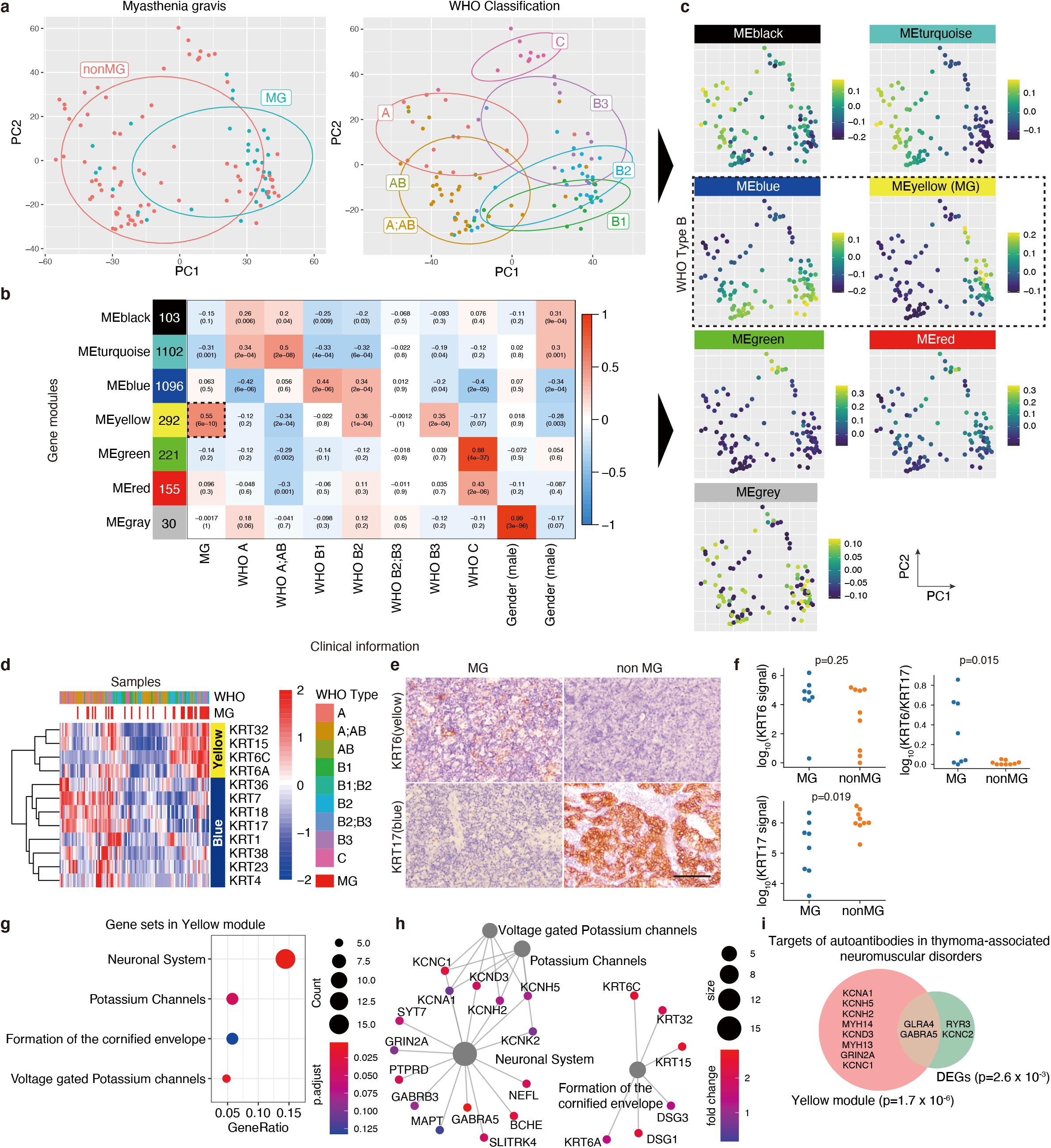
Massive transcriptome profiling of thymoma and MG-specific expression of neuro-related genes. a, PCA plots for transcription profile of thymomas from 116 patients. The left panel shows the disease status, MG or non-MG, and the right panel shows WHO classification based on histology. b, Gene modules defined using WGCNA and the association with MG, WHO classification, gender, and age at diagnosis. Numbers in colored boxes on the left are the number of genes included in each module. The numbers in the heatmap show the correlation (upper) and the *P-value* (lower). c, Eigengenes of each module for each patient on the PCA plot. d, Heatmap of the gene expression of keratins in the yellow and blue modules. The color represents the Z-score of normalized expression by DESeq2. WHO classification and MG status were shown at the top of the heatmap. e, Immunohistochemical (IHC) staining of KRT6 and KRT17 in MG and non-MG-thymoma. The scale bar: 100μm. f, Protein levels of KRT6 and KR17, and KRT6 normalized by KRT17 in MG and non-MG-thymoma quantified using microscopic images (details in Supplementary Data Fig. 1,2 and Methods). The signals were analyzed using a two-sided Mann-Whitney *U* test. g, Significantly enriched REACTOME pathways in the yellow module. The node size represents the number of genes included in each pathway, and the color represents the adjusted *P*-value of the enrichment. The pathways were sorted by the ratio of genes included in the yellow module. h, Genes in enriched REACTOME pathway in the yellow module. Genes with *log*_*2*_ *fold change* > 1 in comparison of MG and non-MG were selected. i, Venn diagram showing overlap of targets of autoantibodies associated with thymoma with genes in the yellow module and upregulated genes in MG. Data were analyzed using a two-sided Fisher’s exact test.

To dissect transcriptome changes unbiasedly, we next adopted the unsupervised gene clustering approach; Weighted Gene Co-expression Network Analysis (WGCNA)^22^ on the large-scale thymoma samples (Extended Data Fig. 2d). We initially constructed gene “modules”, each of which were composed of a set of genes showing correlated gene expression. In the thymoma samples, seven modules consisting of 30-1102 genes were obtained and represented by colors (Supplementary Table 2). We next investigated the association of clinical information with a representative gene expression of each module, calculated as eigengene (Fig. 1b). MG had a most significant correlation with the yellow module (*ρ* = 0.55, *P* = 6 x 10^−10^, Extended Data Fig. 2e) among modules. Each type of the WHO classification corresponded to different modules, respectively; i.e., A, AB - black, turquoise; B1, B2 - blue; B2, B3 - yellow; C - green, red. The grey module was strongly associated with gender, and the black and turquoise modules with age at diagnosis. On the PCA plot based on the transcriptome, the enrichment of each module was well-coordinated with the profile of WHO types, suggesting that the heterogeneity of thymoma can be represented by the gene modules (Fig. 1c). Next, we investigated the detailed gene profiles of the modules associated with MG. In accordance with that MG was particularly enriched in the WHO type B epidemiologically^20^ and in TCGA samples (Extended Data Fig. 2a), the yellow module linked to MG was associated with type B2 and B3 (Fig. 1c). In contrast, the blue module, which was independent of MG, was also associated with types B1 and B2 (Fig. 1c). To distinguish the yellow module from the blue one, we selected cytokeratins, which have various isotypes specific to tissues. Within cytokeratin isotypes, *KRT6A*, *KRT6C*, *KRT15* were specific to the yellow module, whereas *KRT7*, *KRT17*, *KRT18* were to the blue module (Fig. 1d). To confirm the difference histologically, we stained KRT6 and KRT17 proteins in thymoma tissue sections and observed the corresponding staining patterns to MG and non-MG-thymoma, respectively (Fig. 1e,f). We next examined the enriched pathways in the yellow module. Intriguingly, when we examined the enriched pathways in the yellow module, the most significantly enriched pathway was neuronal systems which included GABA receptors (*GABRA5*, *GABRB3*), neurofilaments (*NEFM*, *NEFL*), voltage-gated potassium channels (*KCNC1*, *KCNH2*, *KCNH5*, *KCND3*), and an NMDA receptor (*GRIN2A*) (Fig. 1g,h, Supplementary Table 3). Formation of the cornified envelope including cytokeratins and ion channel transport were also enriched in the yellow module. On the other hand, the other modules did not show the enrichment in neuronal systems but instead showed the enrichment of different types of pathways, such as interleukin-4 and interleukin-13 signaling in the blue module, extracellular matrix organization in the green module, and interleukin-10 signaling in the red module (Extended Data Fig. 2f, Supplementary Table 4).

Thymoma has been shown to associate with paraneoplastic neurological diseases, such as encephalitis and myositis, besides myasthenia gravis^3,23^. We observed the significant overlap of candidate target antigens of thymoma-relating autoantibodies (Supplementary Table 5,6) with the yellow module genes (odds ratio = 7.87, *P* = 1.65 x 10^−6^) and more weakly with the differentially expressed genes between MG and non-MG (odds ratio = 7.42, *P* = 2.67 x 10^−3^). The overlap included an NMDA receptor (*GRIN2*), voltage-gated potassium channels (*KCNH2*, *KCNC1*, *KCNA1*, *KCNC2*, *KCND3*, *KCNH5*), a glycine receptor (*GLRA4*), a GABA receptor (*GABRA5*), and a ryanodine receptor (*RYR3*) (Fig. 1i).

We also examined immune profiles of MG, such as T cell receptor (TCR)/B cell receptor (BCR) diversity and viral infections. The diversity of immunoglobulins in MG-thymoma was lower than that of non-MG-thymoma, suggesting that B-cell maturation and expansion were occurred in MG-thymoma (Extended Data Fig. 3a). The diversity of TCR was mostly unchanged between them, but the composition rate of a TCR alpha chain J, *TRAJ24*, was high in MG (*P*_*adj*_ = 9.6 x 10^−4^; Extended Data Fig. 3b), and especially the *TRAJ24*-*TRAV13-2* combination was 7.50-fold more frequent in MG-thymoma (Extended Data Fig. 3c). To assess the effect of HLA on MG susceptibility, we determined major alleles of the HLA class I and II in MG using the same TCGA bulk RNA-seq dataset. The strongest association was observed in *DQA1*01:04* (odds ratio = 4.43, *P* = 0.050), followed by *DQB1*05:03* (odds ratio = 4.25, *p* = 0.056), *A*24:02* (odds ratio = 2.84, *P* = 0.058, also reported by Machens *et al.,*^24^) though all associations were below the significance level (Extended Data Fig. 3d). MG development has also been shown to associate with viral infections, including SARS-CoV2^25^ and Epstein-Barr virus^26,27^. Therefore, we next examined infected viruses in thymoma using the TCGA dataset. Although various viral transcripts such as Epstein-Barr virus and herpesvirus 6A were detected in MG-thymoma, no significant association with viruses was observed (*False Discovery Rate* (*FDR*) < 0.1, Extended Data Fig. 2e). In addition, we could not find any significant somatic mutations associated with MG, whereas missense mutations in *GTF2I* were observed in 49% of thymoma patients as previously reported^28^ (Extended Data Fig 4). Altogether, the unbiased large-scale omics analysis revealed MG-specific expression of neuromuscular molecules and the distortion in the diversity of TCRs and BCRs.

### Single-cell profiling of thymoma and PBMCs from MG patients

To clarify the source of neurological molecules and the surrounding immune environments in MG-thymoma, we conducted single-cell RNA sequencing (scRNAseq) experiments of thymoma and peripheral blood mononuclear cells (PBMCs) derived from four MG patients (Fig. 2a, Supplementary Fig. 3). The patients consisted of three females and one male, had not received immunosuppressive therapy preoperatively except for one patient, ranged in age from 35 to 55 years, and had thymoma type AB-B2 (Supplementary Table 7). Using a droplet-based single-cell isolation method, we profiled 33,839 cells from thymoma and 30,810 cells from PBMCs and identified 49 clusters upon them (Fig. 2b,c, Extended Data Fig. 5a,b). The cell annotation of PBMCs and the thymus was well-concordant with the previously reported scRNAseq experiments for healthy PBMCs^29^ and the thymus^30^ (Extended Data Fig. 5d,e), and each cluster was well-separated by the specifically expressed genes. (Fig. 2d, Extended Fig. 5c, Supplementary Table 8). In the latter parts, we analyzed the detailed expression profiles of the major clusters; stromal cells, T cells, and B cells, of MG-thymoma.

**Figure 2.**
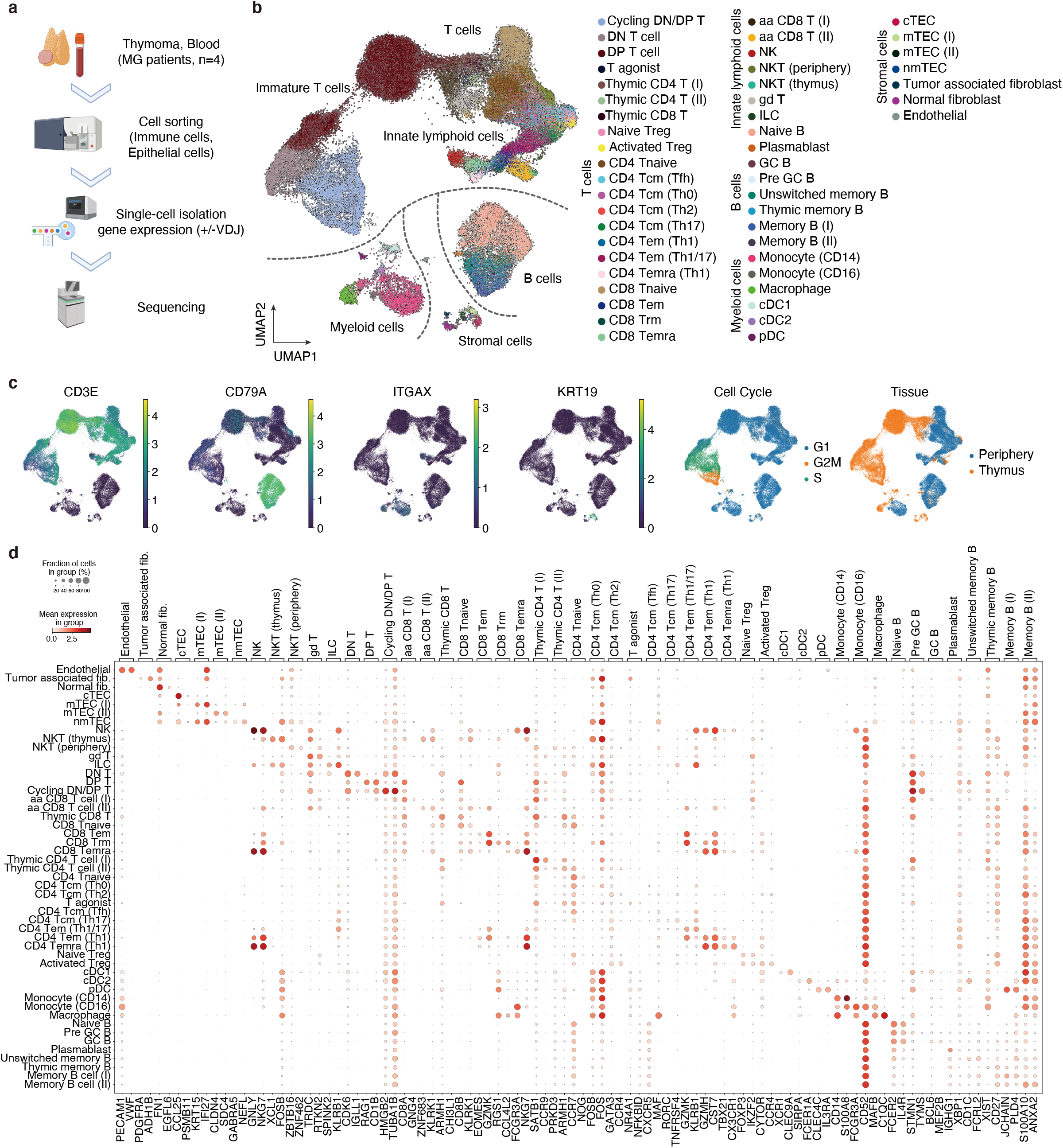
Overview of scRNAseq of thymoma and blood from MG patients. a, The experimental design of scRNAseq. Immune cells and non-immune cells from MG-thymoma and immune cells from the blood of corresponding patients were collected for scRNAseq. b, UMAP plot for 65,935 cells displaying the 49 clusters from thymoma and blood of MG patients. c, UMAP plot of marker genes, inferred cell cycle, and tissue origins. d, Dot plot depicting signature genes’ mean expression levels and percentage of cells expressing them across clusters. The detailed dot plot is shown in Extended Fig. 5c.

### Identification of a unique thymic epithelial cell cluster in MG-thymoma

We profiled stromal cells of thymoma firstly. Clustered stromal cells corresponded to endothelial cells (positive for *PECAM1*/CD31, *VWF*), normal fibroblasts (*FN1*, *EGFL6*), tumor-associated fibroblasts (TAFs; *PDGFRA*, *ADH1B*), and thymic epithelial cells (TECs; *KRT19*, *S100A14*) (Fig. 3a, Extended Data Fig. 6a). We then extracted the TEC cluster and re-clustered them into cTEC (*CCL25*, *PSMB11*) and mTEC (*CCL19*, *KRT7*) clusters (Fig. 3b, c). The mTECs further fell into 3 clusters; mTEC(I) specifically expressing *KRT15* and *IFI27*; mTEC(II) expressing *CLDN4* and *KRT7*; and the unique mTECs expressing neuromuscular-related molecules (Fig. 3d). Cells in this unique mTEC cluster also expressed brain-specific genes included in the yellow module, such as *GABRA5*, *MAP2*, *NEFL*, *NEFM*, *SOX15*, *TF*. Their ectopic expression was also confirmed immunohistochemically in MG-thymoma tissue sections (Fig. 3e, Extended Data Fig. 6b,c). GABRA5 as one of the neuronal molecules expressed in the unique cluster and the cytokeratin KRT6, which belongs to the yellow module, were detected in identical cells (odds ratio = 50.6, *P* < 10^−16^), with the cytoplasm and the pericellular localization of the cells, respectively (Fig. 3g,h, Suoolementary Fig. 4). Due to the atypical expression profile of the cluster, we named the population neuromuscular-mTECs or nmTECs. nmTECs also expressed some of the targets of autoantibodies in thymoma-associated neuromuscular disorders highlighted in TCGA bulk RNA-seq analysis in Fig. 1i (Extended Fig. 6d). To assess the counterparts in the normal thymus, we compared scRNAseq data of thymoma TECs with that of the normal thymus previously published^30^. Thymoma nmTECs were partially correlated with an immature TEC cluster (mcTECs) and not with myoid cells (TEC(myo)) and neuroendocrine cells (TEC(neuro)) in the normal thymus (Fig. 3f). Next, to clarify the biological characteristics of nmTECs, gene set enrichment analysis of nmTECs was performed using the REACTOME gene sets. nmTECs showed the enhancement of pathways such as TP53 activation and pathways in cancer (Fig. 3i,l). nmTECs showed the highest number of detected reads per cell, while markers for other cell-types such as T cells and B cells were not detected in their expression (Fig. 3m, Extended Data Fig. 6e), suggesting that nmTECs are tumorous cells and not doublets. nmTECs also showed the enrichment in E3 ubiquitin ligases, IFNγ signaling, and class I MHC mediated antigen processing and presentation (Fig. 3i-k). Although MHC class II antigen presentation was not significantly enriched in the pathway analysis, nmTECs showed upregulation of HLA class II molecules and IFNγ together with downstream molecules of IFNγ signaling; *STAT1*, *IRF1*, *CIITA*, which have been shown to activate MHC class II regulations^31,31,32^. It suggests that nmTECs may possess a high capability of antigen-presentation via HLA class II (Fig3.n). *AIRE* and *FEZF2*, which have been reported to be involved in the production of self-antigens in mTECs, and tissue-restricted antigens (TRAs) were expressed in a few cells of mTEC(I) cells, but not in nmTECs (Extended Data Fig 6f). These observations thus suggest that nmTECs is a unique population producing neuromuscular-related molecules with active antigen presentation via MHC class I and II molecules.

**Figure 3.**
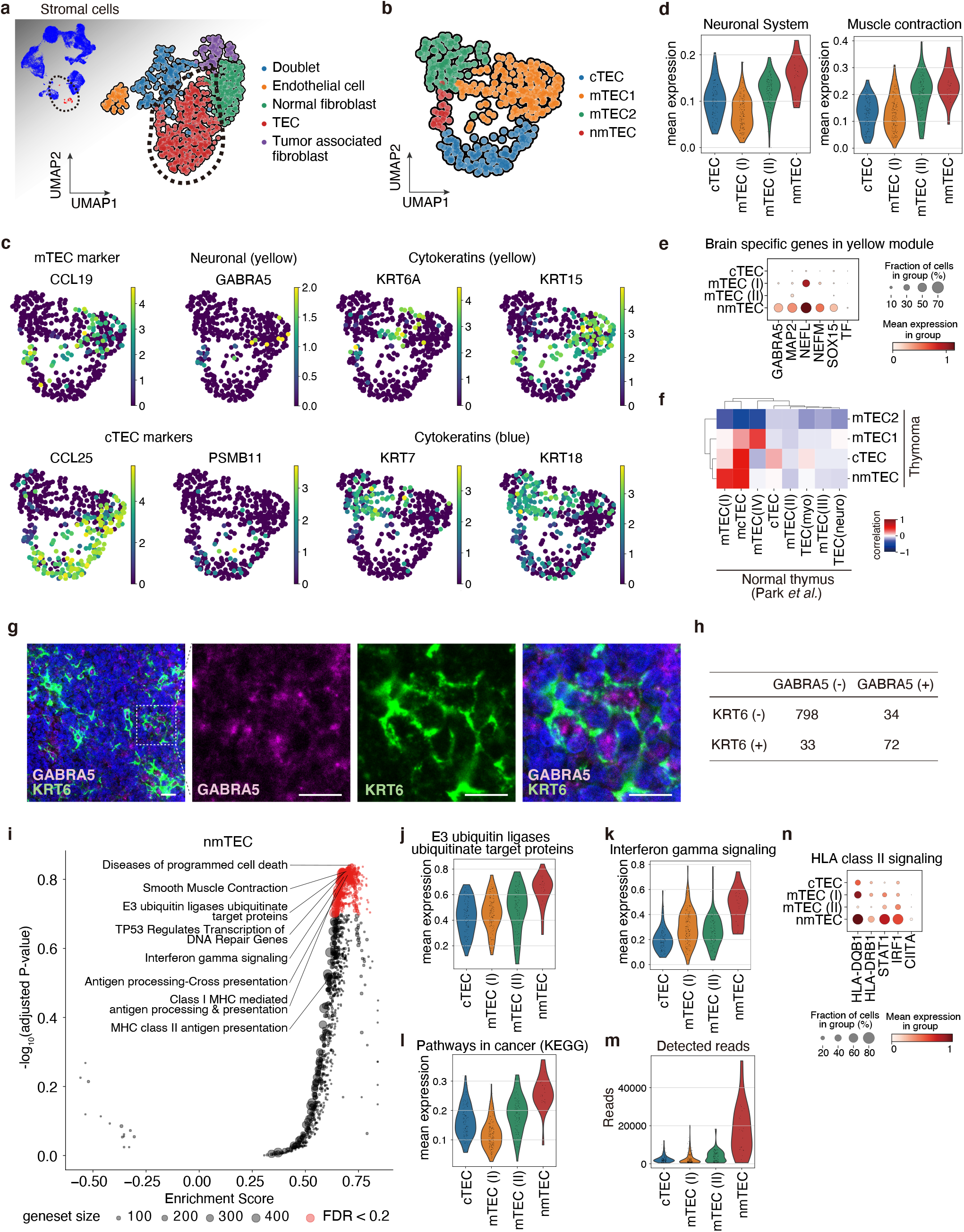
Neuromuscular thymic epithelial cells (nmTECs) expressed neuromuscular genes, IFN gamma signaling pathway genes, and HLA molecules. a,b, UMAP embedding for stromal clusters (a) and thymic epithelial cells (TECs) clusters (b) in thymoma. c, Gene expression of marker genes on UMAP embedding. d, Violin plots of mean expression of the REACTOME gene sets; Neuronal System (left) and Muscle contraction (right) in TEC clusters. e, Dot plot of the yellow module genes. Corresponding protein expressions were also confirmed using IHC (Extended Data Fig. 6b). f, Heatmap showing correlation of transcriptional profile with TEC cells in thymoma (this publication) and a normal thymus (Park *et al.*^30^). g, Immunofluorescence staining for confirming the presence of nmTECs positive for GABRA5 (red), KRT6 (green), and DAPI (blue). Scale bars: 20μm. h, Cross table showing cell numbers of GABRA5 positive/negative and KRT6 positive/negative cells in IHC slides. Data were analyzed using a two-sided Fisher’s exact test. i, Volcano plot showing REACTOME gene sets enriched in nmTECs. j-m, Violin plots of mean expression of the gene sets (j-l) and the number of detected reads per cell (m). j and k represent the significantly enriched REACTOME gene sets and l. represents the KEGG gene set, Pathways in cancer. n, Dot plot of gene expression of HLA class II-related molecules in TECs.

### Dynamics of myeloid cells in MG-thymoma

To explore the MG-specific immune environment in thymoma, we next profiled myeloid cells. We identified six myeloid clusters in thymoma and PBMCs (Extended Data Fig. 7a,b). Monocytes were dominated in PBMCs, while macrophages and dendritic cells were populated mostly in thymoma (Extended Data Fig. 7e). Among clusters, type 2 conventional dendritic cells or cDC2s (*CLEC10A*, *FCER1A*, *ITGAX*/CD11c), which preferentially polarize toward T_H_2, T_H_17, and T_FH_ responses^33,34^, were inferred to migrate from the periphery into thymoma from RNA velocity^35^ (Extended Data Fig. 7c-f).

### B cell maturation with ectopic germinal center formation in MG-thymoma

Since B cells are the source of the autoantibodies causative for MG, we next assessed B cell dynamics in MG-thymoma. To determine the subpopulations of B cells, we categorized them into eight distinct B cell clusters. Notably, we found a population forming a germinal center (GC; positive for *BCL6*, *MEF2B*) in MG-thymoma (Fig. 4a), while GC B cells were not detected in the normal thymus (Extended Data Fig. 5d). The formation of ectopic germinal centers in MG-thymoma was also histologically confirmed by H&E staining (Fig. 4f). Based on the expression of immunoglobulins, B cells were divided into three groups; 1) Naive, GC, pre-GC (*IGHM*, *IGHD*, *IGHG3* high); 2) memory B cells (*IGHA1*, *IGHA2*, *IGHG2* high); and 3) plasmablasts (*IGHG1*, *IGHG3*, *IGHG4* high) (Fig. 4b). We also observed that a pre-GC B cell population (*STMN1*, *TCL1A*) was preferentially enriched in thymoma (Fig. 4c). The RNA velocity analysis showed that pre-GC cells were directed from naive B cells toward GC B cells, memory B cells, and plasmablasts in MG-thymoma (Extended Data Fig. 8a), suggesting that the B cell maturation progresses normally in the MG-thymoma. In addition, Pre-GC, GC, thymic memory B cells, and plasmablasts were enriched in thymoma compared to PBMCs (Fig. 4d,e).

**Figure 4.**
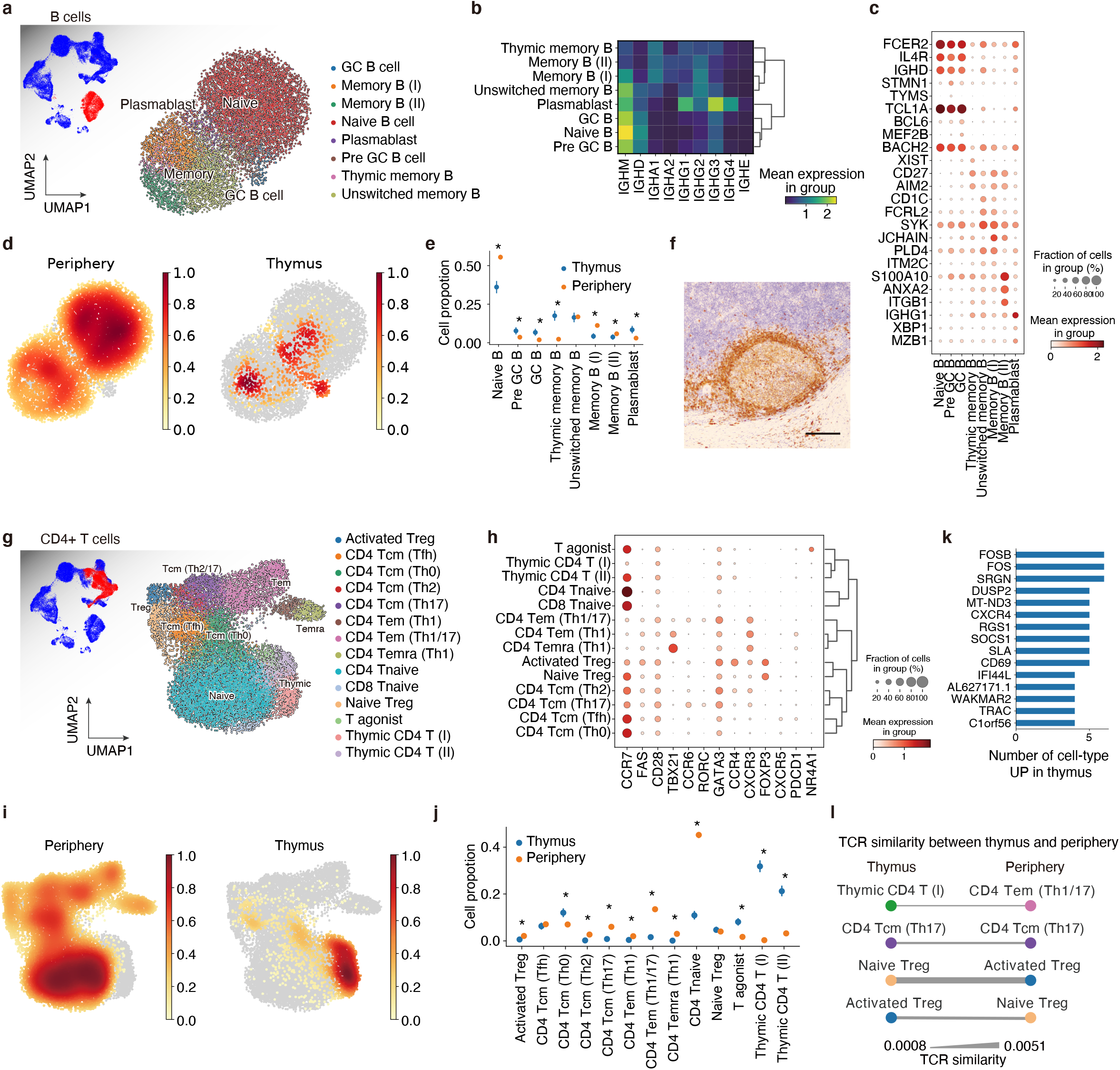
Immune cell landscape elucidates GC formation, T_H_0-T_FH_ enhancement, and Treg recirculation in MG-thymoma. a, UMAP embedding for B cell clusters of thymoma and peripheral blood. b, Heatmap of immunoglobulin expressions in each B cell cluster. Mean expressions in each group are shown as a heatmap. c, Dot plot of gene expression of marker genes of each B cell cluster. d, Density plots showing B cell accumulation in the periphery (left) and thymus (right). e, Cell proportion of each B cell cluster in thymoma and peripheral blood. f, Representative IHC image of germinal center in MG-thymoma stained for CD79A. Scale bar: 100μm. g, UMAP embedding for CD4^+^ T cell clusters of thymoma and peripheral blood. h, Dot plot of gene expression of marker genes of each T cell cluster. i, Density plots showing T cell accumulation in the periphery (left) and thymus (right). j, Cell proportion of each T cell cluster in thymoma and peripheral blood. k, Bar plot of thymus specific genes across CD4^+^ T cell clusters ranked by the number of cell types where each gene was upregulated (*P*_*adj*_ < 0.05 and *log*_*2*_ *fold change* > 1) in mature CD4^+^ T cells. l, TCR similarity between peripheral blood and thymoma. The thicknesses of edges represents TCR similarity. \**FDR* < 0.05 in e and j. Statical procedures in e and j are described in Methods. GC, germinal center.

### T cell polarization in MG-thymoma

In the thymus, T cells are characteristically educated by antigen-presenting cells, including mTECs, and abnormalities of the antigen-presentation frequently associate with autoimmune diseases via T cell dysfunction^36,37^. Therefore, we next investigated whether T cells in thymoma were engaged in MG pathogenesis (Extended Data Fig. 8d). We observed the existence of immature and mature T cells in thymoma, suggesting that the physiological T cell development was maintained even in MG-thymoma. Among populations, we identified a thymoma-specific mature T-cell population, CD8^+^ tissue-resident memory T cell (CD8 T_RM_) expressing *CXCR6* as seen in other tissues such as the lung^38,39^ and skin^40^ (Extended Data Fig. 8d). We next focused on CD4^+^ T cell clusters, which are essential for B cell activation. In thymoma and PBMCs, we identified 13 specific clusters, which corresponded to cells in the process of differentiation, i.e., immature thymic CD4^+^ T cells to terminally differentiated effector memory CD4^+^ T cells (CD4 T_EMRA_). T cells after thymic selection contained CD4^+^ naive T cells (CD4 T_NAIVE_; *CCR7*^+^ *FAS*^−^), CD4^+^ central memory T cells (CD4 T_CM_; *CCR7*^+^ *FAS*^+^), effector memory T cells (CD4 T_EM_; *CCR7*^−^ *FAS*^+^), and terminally differentiated effector memory CD4^+^ T cells (CD4 T_EMRA_; *FAS*^+^ *CD28*^−^) (Fig. 4g,h). We also identified T cell polarizations using characteristic transcription factors and chemokine receptors such as T_H_1 (*TBX21*/Tbet) in T_EM_ and T_EMRA_, T_H_2 (*GATA3*, *CCR4*) in T_CM_, T_H_17 (*RORC*, *CCR6*) in T_CM_, T follicular helper cells (T_FH_; *CXCR5*, *PDCD1*) in T_CM_ (Fig. 4h). These cell annotations were also concordant with the bulk RNA-seq dataset of purified T cells^41^ (Extended Data Fig. 8e). When we assessed the tissue localization of these cells, CD4 T_CM_ (T_H_0) was more abundant in the thymus, and CD4 T_CM_ (T_FH_) was equally abundant in the thymus, whereas other memory T cells such as CD4 T_CM_ (T_H_2), CD4 T_CM_ (T_H_17) were significantly more abundant in the periphery, suggesting that T_H_0-T_FH_ axis are prominent in the thymus (Fig. 4i,j). Next, to infer T cell dynamics between thymoma and periphery, we investigated the commonalities of T-cell receptor (TCR) repertoires of these cell populations. Strong clonal expansions were observed in T_H_1 prone clusters, including CD4 T_EM_ (T_H_1/17), CD4 T_EM_ (T_H_1) and T_EMRA_ (T_H_1), and also slight clonal expansions in CD4 T_CM_ (T_H_2), CD4 T_CM_ (T_H_17), and activated T_reg_ cells (Extended Data Fig. 8f). By examining TCR similarity between the thymus and periphery for each cluster, Treg cells showed higher levels of TCR similarity between the thymus and the periphery, compared to the other cell populations (Fig. 4l). This suggests that naive T_reg_ cells are activated in thymoma aberrantly and circulated into the periphery. In addition, a chemokine receptor *CXCR4* was preferentially expressed in thymic mature T cells (Fig. 4k, Extended Data Fig. 8g). A couple of thymic T cell-specific genes such as *CD69*, *SOCS1* (STAT-Induced STAT Inhibitor 1) and *RGS1* (Regulator Of G-Protein Signaling 1) were also expressed in B cells in thymoma, but not in periphery, suggesting that these genes were regulated by a shared tissue-specific program between T and B cells (Extended Data Fig. 8b,c). Taken together, a detailed analysis of B and T cells revealed that MG-thymoma kept primary lymphoid tissue characteristics for T cell education and gained abnormal inflammatory profiles with ectopic GC formations.

### Cell-cell interaction inference

To analyze the communications among cells, we next inferred cell-cell interaction by integrating single-cell data with a curated ligand-receptor pair database through a bioinformatics application, *CellPhoneDB*^42^ (Fig. 5a). The cell fraction possessing the highest number of intercellular interactions was nmTECs (Fig. 5b). The cell-cell interaction network analysis showed that nmTECs acted as a hub in the network and interacted with myeloid cells, T cells, B cells, tumor-associated fibroblasts, and endothelial cells (Fig. 5c). nmTECs and tumor-associated fibroblasts preferentially expressed *CXCL12*, and thymic B cells and helper T cells, including T_FH_ and T_reg_ cells, expressed its receptor, *CXCR4*. Given that the *CXCR4*-*CXCL12* axis has been shown to play a key role in T cell homing in synovial tissues of rheumatoid arthritis^43^, neurogenesis^44^, and maintenance of hematopoietic stem cells^45^, the interaction may be important for nmTEC-mediated T cell regulation (Fig. 5d). *CXCR5* was expressed in B cells, T_FH_, and CD8 T_RM_, while its ligand, *CXCL13*, was expressed in T_FH_ and CD8 T_RM_, suggesting that the putative role of *CXCR*5-*CXCL13* for T-B interaction in thymoma. This predicted interaction was consistent with the previous findings^46^. We also predicted the interactions of nmTECs with vascular endothelial cells via *VEGFA* and *VEGFE* and with tumor-associated fibroblasts via *PDGFA*-*PDGFRA*. To verify the predicted interaction, we performed immunostaining of CD31 on MG-thymoma sections and observed that GABRA5^+^ nmTECs were in proximity to CD31^+^ vascular endothelial cells (Fig. 5e-g, Supplementary Fig. 5). These observations suggest that nmTECs may promote angiogenesis via the interaction of vascular endothelial cells.

**Figure 5.**
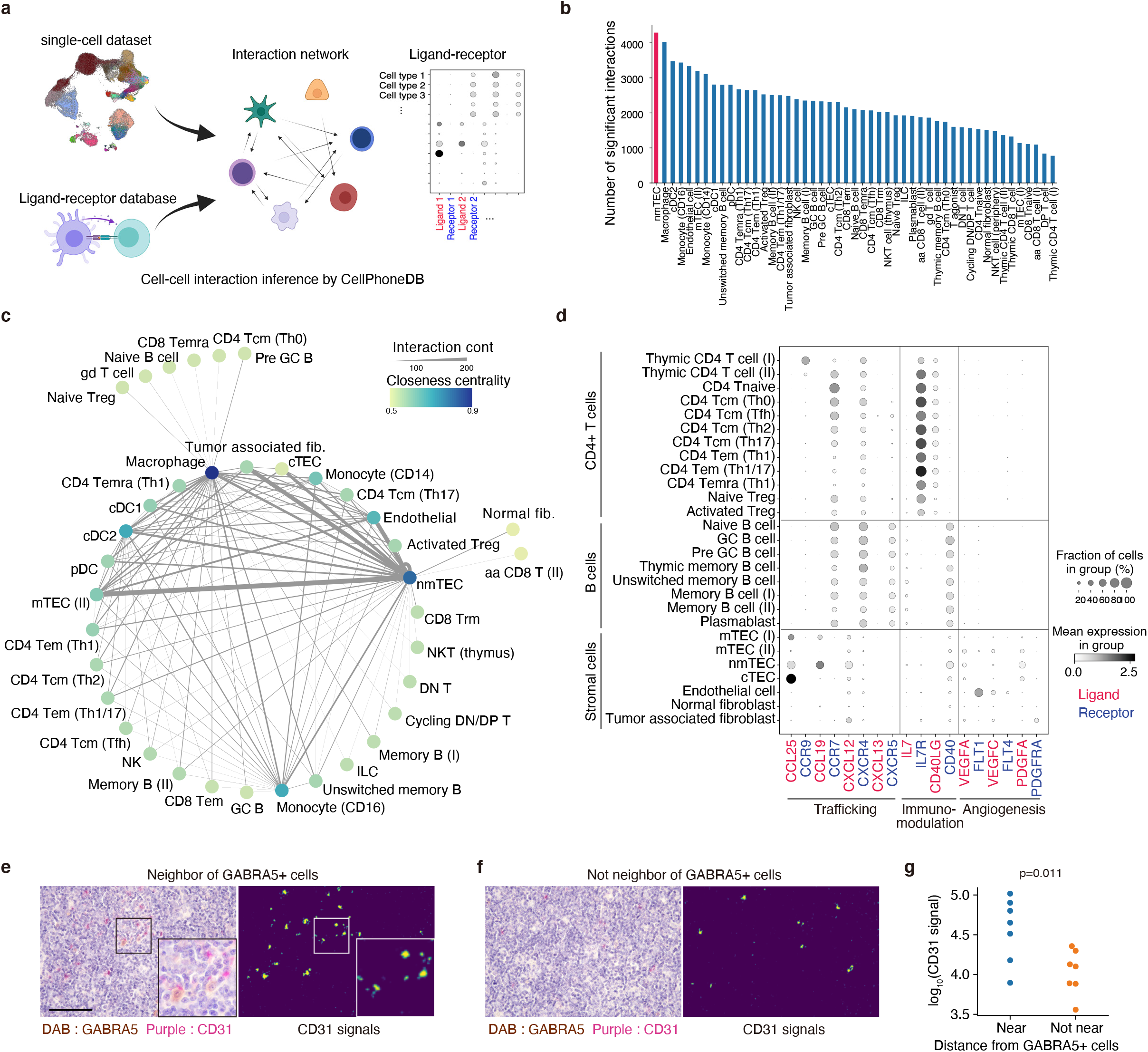
nmTECs strongly associated with epithelial cells, myeloid cells, and T cells with characteristic ligand-receptor pairs. a, Schematic view of the cell-cell interaction analysis. b, Bar chart showing the number of significant interactions with other cell types in each cell type. c, Cell-cell interaction network inferred from scRNAseq data. Each node represents a cell type, and the thickness of each edge represents the number of significant interactions. Edges with less than 75 significant interactions were removed. d, Dot plot of gene expression of ligand-receptor pairs involved in trafficking, immunomodulation, and angiogenesis in CD4^+^ T cells, B cells, and stromal cells. e,f, Representative images of the colocalization of nmTECs (GABRA5; DAB) and endothelial cells (CD31; purple) in the vicinity of GABRA5^+^ cells (e) and not in the vicinity of GABRA5^+^ cells (f). Scale bar: 100μm. g, Protein levels of CD31 near and not near from GABRA5+ cells in MG-thymoma quantified using microscopic images (details in Supplementary Data Fig. 4 and Methods). For each group, seven areas from four MG patients were quantified. The signals were analyzed using a two-sided Mann-Whitney *U* test.

### Integrative analysis of MG pathology across cell types

Recently, a computational method has been developed to infer the cell proportions from bulk RNA-seq datasets using references constituted by scRNAseq^47^. To identify cell populations enriched in MG, we estimated cell distribution by deconvolution of large-scale bulk RNA-seq of thymomas in the TCGA database, using detailed single-cell annotation defined in the previous sections. Among cell populations, cTECs were accumulated in WHO type A; mTECs in type A, B3, C; and immature T cells in type B1 thymoma (Extended Data Fig. 9a). These observations were concordant with the phenotypes defined by WHO classification, suggesting that the deconvolution was functioning well. The numbers of cycling DN/DP T cells and endothelial cells were decreased and increased, respectively, along with age (Extended Data Fig. 9b). The most significantly associated cell population to MG was nmTECs, followed by GC B cells and cDC2s (Fig. 6a-c).

**Figure 6.**
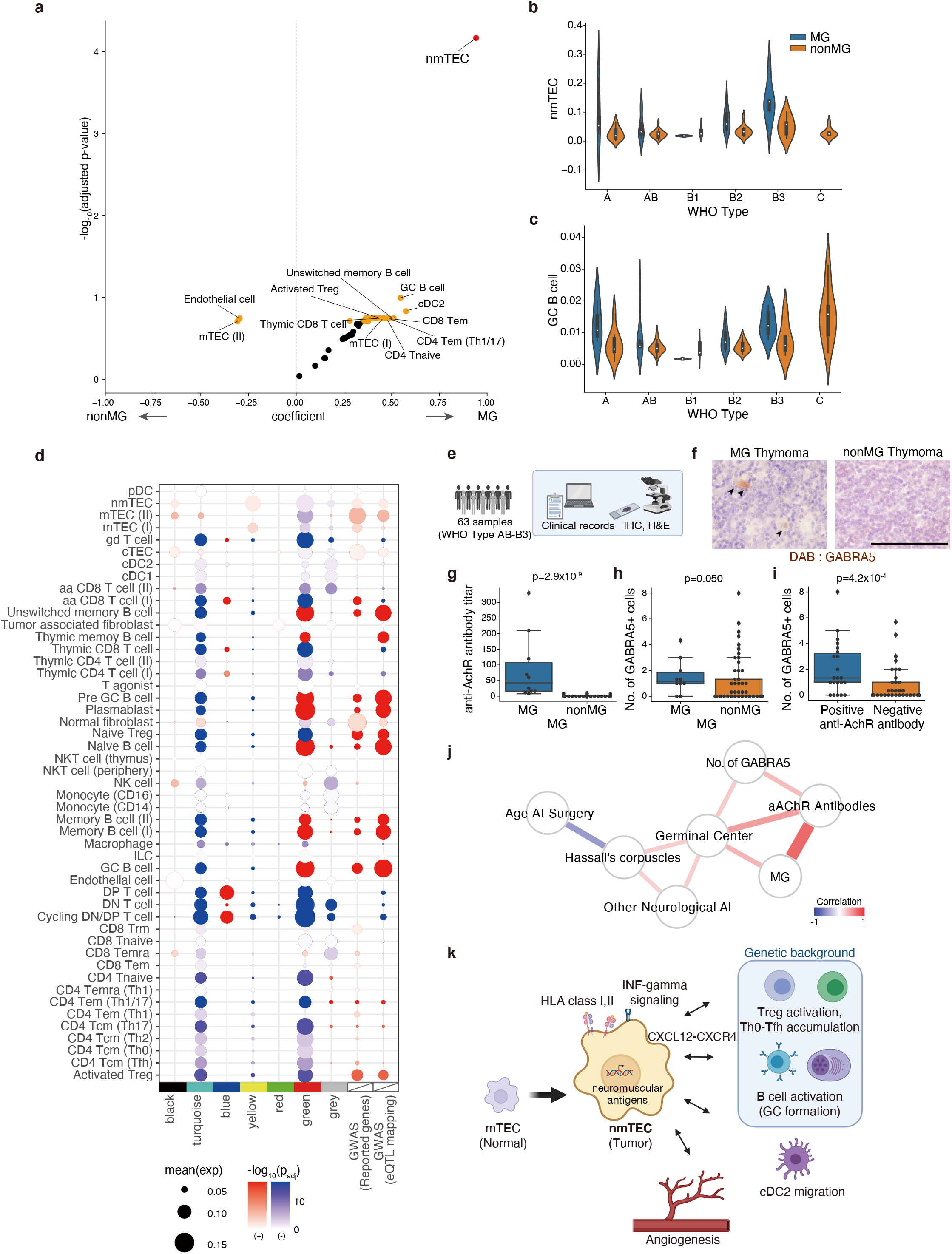
Cell-type wide analysis exhibits nmTECs and GC B cells associate with MG. a, Volcano plot showing the association with MG for deconvoluted cell proportion based on TCGA bulk RNA-seq dataset with the reference defined in our scRNAseq analysis. Red dot *FDR* < 0.05, orange dots *FDR* < 0.2. b,c, Violin plots of the inferred cell proportion for nmTECs (b) and GC B cells (c) partitioned by WHO classification and MG status. d, Expression enrichment of gene modules, targets of autoantibodies in thymoma-associated neuromuscular disorders, and GWAS reported genes for EOMG and LOMG. The enrichment score for each gene set was analyzed using a two-sided Mann-Whitney *U* test across cell-type, and the adjusted *P*-value was calculated. A positive correlation is colored in red, and a negative correlation is in blue. e, Strategy of histological assessment by an independent cohort. f, Representative Immunohistochemical (IHC) staining images of GABRA5 in MG (left) and non-MG (right) thymoma. Arrowheads indicate GABRA5-positive cells. Scale bar: 100 μm. g-i, Box plots of anti-AChR antibody titer (nmol/L) (g) and the number of GABRA5-positive cells in thymoma (h) in MG and non-MG-thymoma patients, and the number of GABRA5-positive cells in thymoma partitioned by anti-AChR antibody titer (i). Data were analyzed using a two-sided Mann-Whitney *U* test. j, Network showing the correlation with clinical and histological features. Anti-AChR antibody titer was tested before the thymectomy. The existence of the germinal center was determined using H&E staining or DAB staining for CD79A. Statistically significant edges with the multiple test correction were retained (*FDR* < 0.2). The edge color represents Pearson’s correlation, and the thickness of the edge represents −*log*_*10*_*FDR*. k, Proposed MG pathology in thymoma. EOMG, early-onset MG; LOMG, late-onset MG.

Next, we examined the contribution of each cell type to each module defined by WGCNA. nmTECs were the most significantly contributed to the yellow module, which was associated with MG (mean expression = 0.092, *P*_*adj*_ < 10^−13^; Fig. 6d, Supplementary Table 11). The blue module, which was also associated with WHO type B, was found to be associated with cycling DN/DP T cells and DP cells (mean expression = 0.092 and = 0.10, *P*_*adj*_ < 10^−100^ and < 10^−100^; Fig. 6d, Supplementary Table 11). The target molecules of thymoma-associated autoantibodies were also enriched in nmTECs significantly (mean expression = 0.014, *P*_*adj*_ =1.4 x 10^−3^; Extended Data Fig. 10c, Supplementary Table 10). To measure genetical effects on each cell population, we listed up myasthenia gravis associated genes reported in three genome-wide association studies (Seldin *et al.*^48^, 532 cases, 2128 controls; Renton *et al.*^49^, 1455 cases, 2465 controls; Gregersen *et al.*^50^, 649 cases, 2596 controls, Supplementary Table 10). We excluded HLA genes, which possessed the most significant signals, to avoid the ambiguity derived from the complex linkage disequilibrium (LD) structure in the HLA regions. We also extracted genes associated with GWAS SNPs in consideration of expression quantitative trait locus (eQTL) and LD structures (see Methods). MG-associated genes in both lists were significantly associated with T_reg_ cells and B cells, including GC B cells and plasmablasts (Fig. 6d, Extended Data Fig. 10a,b, Supplementary Table 11). Overall, these analyses indicated that nmTECs, GC B cells, and cDC2s were atypically increased in MG-thymoma and that the genetic effects associated with MG were mainly accumulated in T and B cells.

### Histological validation of the MG-associated phenotypes

To validate the MG-associated changes in another cohort, we examined tissue specimens from 63 WHO type AB-B3 thymoma surgery cases with the clinical records (Fig. 6e). To quantify the amount of nmTECs, we performed immunostaining for GABRA5 (Fig. 6f) and found that the number of GABRA5 positive cells was higher in MG (*P* = 0.050) and more significantly in anti-AChR antibody-positive thymoma patients (*P* = 4.2 x 10^−4^, Fig. 6g-i). In addition, the presence of germinal centers determined by H&E staining was associated with the increase of anti-AChR antibodies, the presence of MG/other neuro-related autoimmune diseases, and the number of GABRA5 positive cells (Fig. 6j). These observations depicted that the emergence of nmTECs was involved in MG pathogenesis in thymoma together with the altered immune cell populations.

## Discussion

In this study, we revealed the pathogenic changes responsible for MG in thymoma by exploring MG-deviated expression at the single-cell level. As a key finding, we identified abnormal expression of neuromuscular molecules specific to MG cases within thymoma. Single-cell RNA-seq and immunohistological examination of MG-thymoma specimens revealed that these neuromuscular expressions were limited in a subpopulation of mTECs (GABRA5^+^KRT6^+^), termed nmTECs. In addition, MG-thymoma developed atypical immune microenvironments with GC formation, B cell maturation, and ectopic neuromuscular expression on nmTECs, providing a holistic picture of the cell dynamics for producing autoantibodies, which was previously known only in fragments (fig. 6k).

While TAMG is caused by autoantibodies against acetylcholine receptors expressed at the neuromuscular junction under normal conditions, the mechanisms by which those autoantibodies are generated have not been clarified so far. In this study, integrated omics analysis showed that responsible antigen-presenting cells to present acetylcholine receptors would be nmTECs in the thymus. mTECs originally possess the ability to express systemic antigens ectopically using a transcription factor, *AIRE,* to eliminate self-reactive T cells^7^. In fact, it has been shown that mTECs acquire a variety of cell polarities such as tuft, keratinocyte-like, and neuroendocrine after *AIRE* expression^51,52^. Therefore, it seems likely that acetylcholine receptor expression by nmTECs would be caused by the intrinsic ability of mTECs to present self-antigens under the negative selection. The expression of autoantigens is also known to be enhanced by IFN-γ^53^. We observed that the IFN-γ signaling cascade in nmTECs was more active than those in normal mTECs, indicating that they present antigens to immune cells more efficiently. Thus, nmTECs would feed self-antigens to autoreactive lymphocytes and trigger pathological GC formation in the thymoma. This also gives rise to the possibility that the physiological production of self-antigens by mTECs might have a risk of inducing autoimmunity. Interestingly, MG-thymoma expresses not only acetylcholine receptors but also various neuromuscular-related antigens associated with other autoimmune diseases, suggesting that the abnormal expression of neuromuscular antigens by nmTECs is also associated with thymoma-associated neuromuscular autoimmune diseases. This may provide clues to elucidate the pathogenesis of a wide range of neurological autoimmune diseases.

We have succeeded in capturing the entire picture of the thymic microenvironment for producing autoantibodies causative for MG. It is widely accepted that mature B cells in the thymus serve as a source of autoantibodies^54^. In addition, GC formation and an increase of Tfh cells in the thymus have been reported as immune changes in MG-thymoma^17,18^. Our results were fully consistent with those observations and further revealed the accumulation of cDC2, which are considered as migrating DCs from the peripherally for supporting B cell maturation^55^, in MG-thymoma. Cell-cell interaction analysis also predicted that the *CXCR4*-*CXCL12*-mediated interaction between lymphocytes and nmTEC in the thymus is one of the key interactions for producing autoantibodies in the thymoma microenvironments. The interaction of nmTECs and lymphocytes together with cDC2, Tfh, GC accumulation suggests that there may be MG-specific immune microenvironments that support the maturation of autoantibody-producing B cells and their migration to the periphery. It has also been reported that anti-AChR antibody-producing cells reside in the bone marrow^56^ and lymph nodes^57^ outside the thymus and that antibody-producing cells continued to circulate in the periphery after thymectomy^58^. The circulation between the thymoma and the periphery also seems to be present in T cells, as suggested by our TCR repertoire analysis. We thus now have a better understanding of the thymoma microenvironment in which autoreactive B cells are maturated with the help of neuromuscular molecule-presenting nmTECs, the construction of GC formation, enhanced Tfh cell activity, and cDC2 accumulation.

One of the remaining questions is whether the expression of neuromuscular molecules by mTECs triggers the MG development. Our data showed that some patients with high expression of neuromuscular genes did not develop MG. Histological analysis also showed that GABRA5-positive, or nmTEC marker-positive, cells were present in some acetylcholine receptor antibody-negative patients. These results suggest that the accumulation of neuromuscular-related antigens induces a pre-disease state and is not a sufficient condition for MG pathogenesis. In other words, MG pathogenesis requires additional factors except for the differentiation of nmTECs. One of the candidate factors is viral infections since viral infections have been reported to be involved in many autoimmune diseases, including MG^25,26,27^, via inducing immune disruption. While we could not detect any virus that significantly correlated with MG, its effect might contribute to the MG pathogenesis as reported. Another pathological factor is the genetic factors. The integrated analysis with GWAS reaffirmed the importance of T cells including T_reg_ cells and B cells as a genetic predisposition for MG pathogenesis. Therefore, MG would be cooperatively developed by the expression of neuromuscular-related antigens, skewed immune microenvironment, genetic backgrounds, and environmental factors including virus infections. Further analysis will be required for addressing the stepwise development of MG.

Finally, we revealed the complex relationship between MG and thymoma from a view of cell composition and the source of neuromuscular molecules causative for MG. We hope that this study will provide useful information for the development of MG therapy.

## Methods

### Human samples

The study using human samples was reviewed and approved by the Research Ethics Committee of Osaka University and carried out in accordance with the guidelines and regulations. Human samples were collected under approved Osaka University’s review board protocols: ID 10038-9 and ID 850-2. Written informed consent was obtained from all donors.

### Immunohistochemistry

All tissue samples were fixed in 10% formalin, embedded in paraffin, cut into 4-μm-thick sections. For DAB staining, Immunohistochemical staining was performed using the Roche BenchMark ULTRA IHC/ISH Staining Module (Ventana Medical Systems) with the Ultra CC1 mild protocol. For Double stains, we performed a second stain for slides that were DAB stained by the Ultra CC1 mild protocol using the Stayright Purple kit (AAT Bioquest). For multicolor fluorescent staining, we stained slides using Opal 4-Color IHC Kits (AKOYA Biosciences) and observed using the Zeiss LSM 710 or LSM 880 confocal microscope and ZEN microscope software (Carl Zeiss). The primary antigens and dilution ratios used are presented in Supplemental Table 12. The scoring of immunohistochemical staining images was supervised by the pathologists (K.K. and S.N.).

### Histological quantifications

For DAB signal quantification, the region with the strongest DAB signal in each slide was captured. Double-stained slides of CD31 and GABRA5 were captured up to two GABRA5-positive areas and an equal number of negative areas from each MG-thymoma specimen under 40x objective. After adjusting the white balance, signals of hematoxylin and DAB or hematoxylin, DAB, and Purple were separated using the reb2hed function in a python package scikit-image (v0.18.1) and quantified the areas above the threshold (Supplementary Fig. 1,2,5). For the distance between GABRA5 signals and KRT6 signals, the distances between the nearest blobs of GABRA5 and KRT6 were measured (Supplementary Fig. 4) were measured. Blobs were defined using the blob_log function provided by a python package scikit-image (0.18.1). The number of nmTECs in Figure 6f was calculated by averaging the number of positive cells in the three regions with the highest accumulation of positive cells under x40 objective using DAB staining for GABRA5. The detection of germinal centers was judged by H&E staining or IHC of CD79A. The existence of Hassall’s corpuscles was judged by HE staining.

### Cell preparation and sequencing of scRNAseq

To ensure the quality of the library, the library preparation of all thymoma and peripheral blood samples was completed by the next day after the collection. Immune cells and thymic epithelial cells were isolated from thymic tissue dissected surgically, as previously described^59^. Briefly, thymic tissue was mechanically disrupted, and the fraction containing lymphocytes was collected. Extracted cells were stained with 7-AAD (BD Biosciences), and live cells were collected as a lymphocyte fraction. The remaining thymic tissue was subjected to enzymatic treatment (Collagenase A (Worthington), DNAse I (Roche, Basel Switzerland), Trypsin/EDTA (nacalai tesque)) and the resulting cells were then subjected to a percoll density gradient centrifugation for the enrichment of thymic epithelial cells. Cells derived from low-density fraction were stained using FITC-labeled anti-EpCAM mAb (dilution: 1/10, HEA-125, Miltenyi Biotec), PE-labeled anti-CD45 mAb (dilution: 1/100, HI30, Biolegend). Dead cells were excluded by 7-AAD staining, and CD45 (low) EpCAM (high) was defined as thymic epithelial cells. Immune cells and thymic epithelial cells were isolated using BD Biosciences FACS Aria II. The gating strategy is described in Supplementary Data Fig. 3. For CD4^+^ T cells and B cells, we first collected PBMCs using Ficoll-Paque (Cytiva). Isolated PBMCs were washed, blocked Fc receptors using Fc Receptor Binding Inhibitor Polyclonal Antibody, Functional Grade, eBioscience™ (Thermo Fisher Scientific), and stained using FITC-labeled anti-CD3 mAb (dilution: 1/100, UCHT1, BD Bioscience), APC-labeled anti-CD4 mAb (dilution: 1/100, RPA-T4, Thermo Fisher Scientific), PE-labeled anti-CD19 mAb (HIB19, BioLegend), Live/Dead (Thermo Fisher Scientific). Then, live-CD3^+^CD4^+^CD19^−^ cells and live-CD3^−^CD4^−^CD19^+^ cells were isolated using BD Biosciences FACS Aria II.

The sorted cells were loaded to Chromium Next GEM Chip G (10x Genomics) on Chromium Controller (10x Genomics) for barcoding and cDNA synthesis. Amplification of the cDNA and the library construction was performed using Chromium Next GEM Single Cell 3’ GEM, Library & Gel Bead Kit v3.1 or Chromium Next GEM Single Cell 3’ Kit v3.1 (10x Genomics) for 3’ profiling and Chromium Next GEM Single Cell 5’ Kit v2 and Chromium Single Cell Human BCR Amplification Kit or Chromium Single Cell Human TCR Amplification Kit (10x Genomics) for 5’ and VDJ profiling according to the manufacturer’s protocol. The libraries were sequenced on NovaSeq6000 (Illumina).

### TCGA-THYM bulk RNA-seq analysis

RNA-seq fastq files for thymoma were downloaded from the GDC Data Portal using gdc-client. Gene expression matrix quantified by HTSeq and clinical information was downloaded through an R package TCGAbiolinks. The detection of differentially expressed genes was performed by DESeq2^21^ (1.30.1) with the design ~ primary_pathology_history_myasthenia_gravis after the removal of mean count below 5. For the visualization of a volcano plot, the lfcShrink function in DESeq2 was applied. Visualizations were performed by the plotPCA function in DESeq2, and R packages EnhancedVolcano, pheatmap, and ggplot2.

### WGCNA analysis

A transformed matrix by the vst function in DESeq2 was used for WGCNA analysis. The top 3000 genes in the variance of the vsd matrix were selected. Then, we calculated the adjacency using the adjacency function with power=5, created Topological Overlap Matrix by TOMsimilarity, calculated the gene tree by hclust against 1 - TOM with method = “average”, and conducted a dynamic tree cut with the following parameters; deepSplit = 2, pamRespectsDendro = FALSE, minClusterSize = 50. The eigengenes of each module were used for the correlation with clinical information. A pathway enrichment analysis was performed utilizing R packages clusterProfiler and ReactiomePA. Genes included in each module or included in the yellow module and with *log*_*2*_ *fold change* > 1, and genes of each module were analyzed using the enrichPathway function.

### Immunoreceptors quantification

The determination and quantification of TCR and BCR were performed by the MiXCR^60^ (v3.0.3) analyze shotgun command with the options; --species hs --starting-material rna --only-productive. The Gini index for CDR3 amino acid sequences was calculated by an in-house program implemented in Python.

For HLA genotyping and quantification, we first aligned fastq reads on the hg38 reference genome using STAR (v2.7.2a). Then, HLA genotypes and expressions were extracted using arcasHLA^61^ (v0.2.0) with IMGT.HLA database (3.24.0) with default parameters.

### Comprehensive virus detection from bulk RNA-seq

The comprehensive viral quantification of RNA-seq was performed by a bioinformatics pipeline; VIRTUS^62^ (v1.2.1), which was composed of fastp, STAR, and Salmon. First, we created indices using createindex.cwl with references downloaded from Gencode v33. Then, we quantified viruses using VIRTUS.PE.cwl with options --hit_cutoff 0 --kz_threshold 0.3.

### Somatic mutation analysis of TCGA-THYM

Mutation data of thymoma was downloaded using an R package, TCGAbiolinks (2.16.4). Visualization was performed by the oncoplot function implemented in an R package, mafttools.

### Bioinformatics analysis of scRNAseq

Sequenced reads were quantified by Cell Ranger (v5.0.0) with pre-built reference refdata-gex-GRCh38-2020-A downloaded at 10x GENOMICS’ website. Quantified expressions were preprocessed and visualized using Scanpy^63^ 1.7.2 and python 3.8.0. IGKV, IGLV, IGHV, IGLC, TRAV, and TRBV genes were removed for the clustering and embedding for the removal of the effect of clonal expansion. Cells with mitochondrial genes were higher than 20%, or detected genes less than 200 were filtered out, then preprocessed by sc.pp.normalize_per_cell with counts_per_cell_after=1e4, sc.pp.log1pp, retained highly variable genes, scaled using sc.tl.scale, and computed principal components using sc.tl.pca. The batch effect of samples was removed by the BBKNN^64^ algorithm. Cells were embedded by UMAP using sc.tl.umap, clustered using sc.tl.leiden, and manually annotated. T cells, B cells, myeloid cells, and stromal cells were extracted, re-clustered from raw counts, and annotated manually through the same procedure where parameters were determined heuristically. The inference of the cell cycle was performed using the sc.tl.score_genes_cell_cycle function following the tutorial (https://nbviewer.jupyter.org/github/theislab/scanpy_usage/blob/master/180209_cell_cycle/cell_cycle.ipynb). The enrichment scores of gene sets such as GWAS reported genes and WGCNA module genes were calculated by the sc.tl.score_genes function of Scanpy. A two-sided Mann-Whitney *U* test was performed for scores of a cluster and that of others by the scipy.stats function. *P-value* correction for multiple tests was conducted using the statmodels package. Gene set enrichment analysis for clusters was performed by prerank test implemented in gseapy with scores calculated by sc.tl.rank_genes_groups with the option method=t-test_overestim_var.

To examine the difference in cell distribution in the thymus and blood, we used Bayesian estimation in consideration of the imbalance in the number of observed cells in the samples. For each cluster, we inferred the difference between *p*_*thymus*_ and *p*_*blood*_ using the following model;

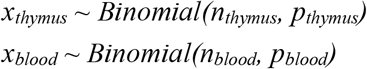

where,

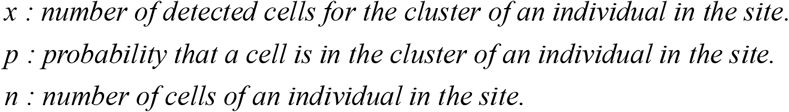

The following was used as a prior distribution of *p*.

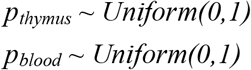

The inference was conducted using a python package pymc3 (3.11.2) using 4 independent chains, 1,000 tuning iterations, and 25,000 additional iterations per chain. Trace plots and R_hat were used to assess the convergence.

For the extraction of thymus specific genes in CD4^+^ T cells (Fig. 4i), we examined the following cell types; CD4 T_NAIVE_, Naive T_reg_, Activated T_reg_, CD4 T_CM_ (T_H_0), CD4 T_CM_ (T_H_17), CD4 T_CM_ (T_FH_), CD4 T_CM_ (T_H_2), CD4 T_EM_ (Th1), CD4 T_EM_ (T_H_1/17), CD4 T_EMRA_ (T_H_1). Similarly, we used the following cell types in B cells (Extended Data Fig. 8c); Naive B cell, Plasmablast, Pre GC B cell, GC B cell, Memory B cell (I), Memory B cell (II), Thymic memory B cell, Unswitched memory B cell.

For the determination of RNA velocity, velocyto run10x was performed with the repeat file hg38_rmsk.gtf downloaded at the UCSC website. The projection of velocities was performed by scVelo (0.2.3)^65^ following the same procedures and parameters as the official tutorial (https://scvelo.readthedocs.io/VelocityBasics/). Visualization was performed by functions of python packages; Scanpy, plotly (4.14.3), matplotlib (3.4.1), and seaborn (0.11.1). Details were described in codes deposited in the Github repository.

### Integration with public single-cell data

H5ad files of scRNAseq data previously reported were downloaded respectively (PBMC: https://atlas.fredhutch.org/data/nygc/multimodal/pbmc_multimodal.h5seurat; normal thymus: 10.5281/zenodo.3711134). Data integration with public single-cell data was performed by the sc.tl.ingest function in Scanpy. For each reference dataset, we extracted highly variable genes, normalized and scaled the expression, and ingested our dataset to reference datasets similarly to our dataset. For the integration of TEC cells and B cells, we re-clustered cells from two datasets with the batch correction instead of the ingestion because some clusters were expected not to have their counterparts leading to the failure of the appropriate ingestion. We first concatenated our data with data from Park et al., removed the batch effect with BBKNN^64^, and calculated correlations using sc.tl.dendrogram because TEC cells in thymoma were expected to be consist of a different set of cells from that in the normal thymus.

### Definition of tissue-restricted antigens (TRA)

To define the list of tissue-restricted antigens (TRA), we used bulk RNA-seq data across tissues provided by Genotype-Tissue Expression (GTEx) project (https://storage.googleapis.com/gtex_analysis_v8/rna_seq_data/GTEx_Analysis_2017-06-05_v8_RNASeQCv1.1.9_gene_tpm.gct.gz). We calculated the Gini index for mean TPM across tissues and extracted genes with Gini index > 0.8 and mean TPM at the maximum expressed tissue > 100 as TRAs (Supplementary Table 9).

### Inference of cell-cell interaction

Cell-cell interaction was inferred by CellPhoneDB (2.1.7), which utilizes abundantly curated ligand-receptor pairs to measure the interactions within single-cell datasets. Statistical test was performed with the default parameters. Dot plots of ligand-receptor pairs were plotted by the cellphondb plot dot_plot function.

### Deconvolution of bulk RNAseq

A deep-learning-based deconvolution tool, Scaden^47^ (v1.1.0) was used for the deconvolution of bulk RNAseq datasets by TCGA. First, we created 30000 simulation datasets with scaden simulate by scaden simulate with option -n 30000. Second, count matrices of our single-cell dataset and TCGA thymoma dataset quantified by HTseq downloaded by TCGAbilkinks were pre-processed by the scaden process command. Then, trained a network by the command scaden train with the option --steps 5000. Lastly, the bulk RNAseq matrix was deconvoluted by scaden predict. Deconvoluted cell proportion was tested using a multiple linear regression provided as the formula.api.ols function by a python package statsmodels (0.12.0) with a model, cells ~ MG + WHO + days_to_birth + Gender + 1.

### Curation of GWAS reported genes

We listed up GWAS-reported genes (*P* < 5×10^−6^) from previous reports^48–50^. Genes in HLA regions were excluded from the list. For eQTL and LD-aware gene mapping, LDexpress (https://ldlink.nci.nih.gov/?tab=ldexpress) in the LDlink suite^66^ was used. We used all populations for the LD reference and all tissues from GTEx v8^67^ for eQTL reference with the threshold, *R*^2^ ≥ 0.1, *P* < 0.1. For each reported locus, we selected a gene that possesses the smallest *P-value*.

### Statistical analysis

All statistical analyses were performed in R (4.0.3) and python (3.8.0). FDR was obtained by the Benjamini-Hochberg procedure implemented by a python package statsmodels (0.12.0). Pearson’s correlation used for Figure 6f was calculated using a python package pingouin (0.3.8). The visualization of a network was performed using Cytoscape (3.8.0)^68^. All other statistical analyses are detailed in the respective sections of the article.

## Code and data availability

TCGA data is available on dbGaP accession phs000178. All source codes will be deposited in the GitHub repository. Single-cell data will be deposited in SingleCellPortal upon the acceptance. Sequence data for single-cell analysis will be deposited in NBDC database upon the acceptance.

## Acknowledgments

We thank Y. Tachibana, K. Funakoshi, and A. Harada for supporting the annotation of single-cell data. We thank T. Sawamura, M. Nihei at the Department of Pathology, Osaka University for supporting the preparation of slides for histological assessments. We thank M. Okumura for his critical advice on the study. This study was supported by the Center for Medical Research and Education, Graduate School of Medicine, Osaka University. We acknowledge the NGS core facility of the Genome Information Research Center at the Research Institute for Microbial Diseases of Osaka University for the support in RNA sequencing. Some illustrations were generated with BioRender.com. The results published here are in part based upon data generated by the TCGA Research Network: https://www.cancer.gov/tcga. This work was supported by Grants-in-Aid by Japanese Society for the Promotion of Science (JSPS) for Specially Promoted Research 16H06295 to S.S., by the Core Research for Evolutional Science and Technology (CREST, no. 17 gm0410016h0006) program from the Japan Science and Technology Agency to S.S. and by Leading Advanced Projects for medical innovation (LEAP, no. 18 gm0010005h0001) from Japan's Agency for Medical Research and Development (AMED) to S.S.

## Author information

### Affiliations

Department of Neurology, Graduate School of Medicine, Osaka University, Suita, Osaka, Japan

Yoshiaki Yasumizu, Hisashi Murata, Makoto Kinoshita, Tatsusada Okuno, Eriko Takeuchi & Hideki Mochizuki

Department of Experimental Immunology, Immunology Frontier Research Center, Osaka University, Suita, Osaka, Japan

Yoshiaki Yasumizu, Yamami Nakamura, Masaya Arai, Yusuke Takeshima, Naganari Ohkura & Shimon Sakaguchi

Integrated Frontier Research for Medical Science Division, Institute for Open and Transdisciplinary Research Initiatives (OTRI), Osaka University, Suita, Osaka, Japan

Yoshiaki Yasumizu, Daisuke Okuzaki & Hideki Mochizuki

Department of Pathology, Graduate School of Medicine, Osaka University, Suita, Osaka, Japan

Satoshi Nojima, Kansuke Kido, Masaharu Kohara & Eiichi Morii

Department of General Thoracic Surgery, Graduate School of Medicine, Osaka University, Suita, Osaka, Japan

Yasushi Shintani & Soichiro Funaki

Genome Information Research Center, Research Institute for Microbial Diseases, Osaka University, Suita, Osaka, Japan

Daisuke Motooka & Daisuke Okuzaki

Department of General Thoracic Surgery, National Hospital Organization Toneyama Hospital, Osaka, Japan

Meinoshin Okumura

## Contributions

Y.Y., T.O., N.O., and H.M. designed all experiments; Y.Y., M.H., K.K., M.Kohara, Y.N., and M.A performed experiments under the supervision of T.O., M.Kinoshita, S.N. and N.O; Y.Y., E.T., S.Suganami, and Y.T. performed bioinformatics analysis; Y.Y. and K.K. diagnosed thymoma pathology under the supervision of S.N. and E.M.; Y.S. and S.F. collected samples for analysis; D.M. and D.O. performed library construction and sequencing; Y.Y., E.T. and M.H. prepared the figures; Y.Y. and N.O. drafted the manuscript; O.T., H.M., E.M., N.O. and S.Sakaguchi supervised the study; S.T. and O.M. provided expert guidance on the manuscript; All authors critically reviewed and edited the final version of the manuscript.

## Extended figure legends

**Extended Data Fig. 1.**
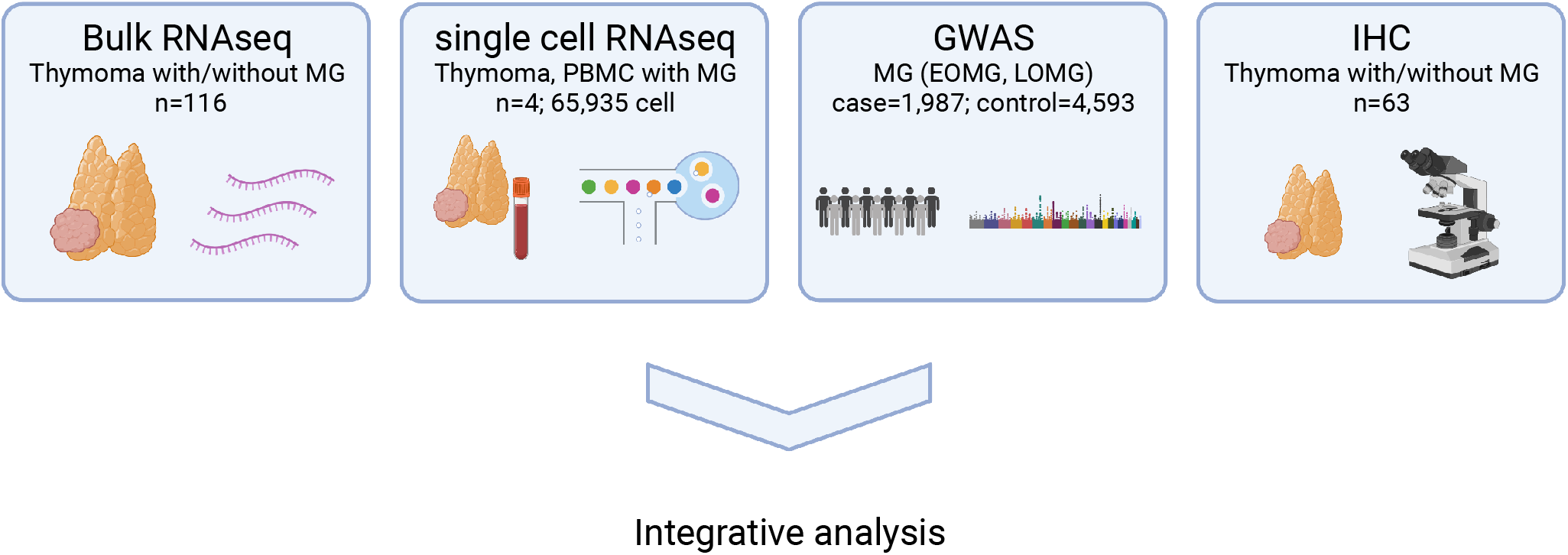
Schematic view of the analysis flow.

**Extended Data Fig. 2.**
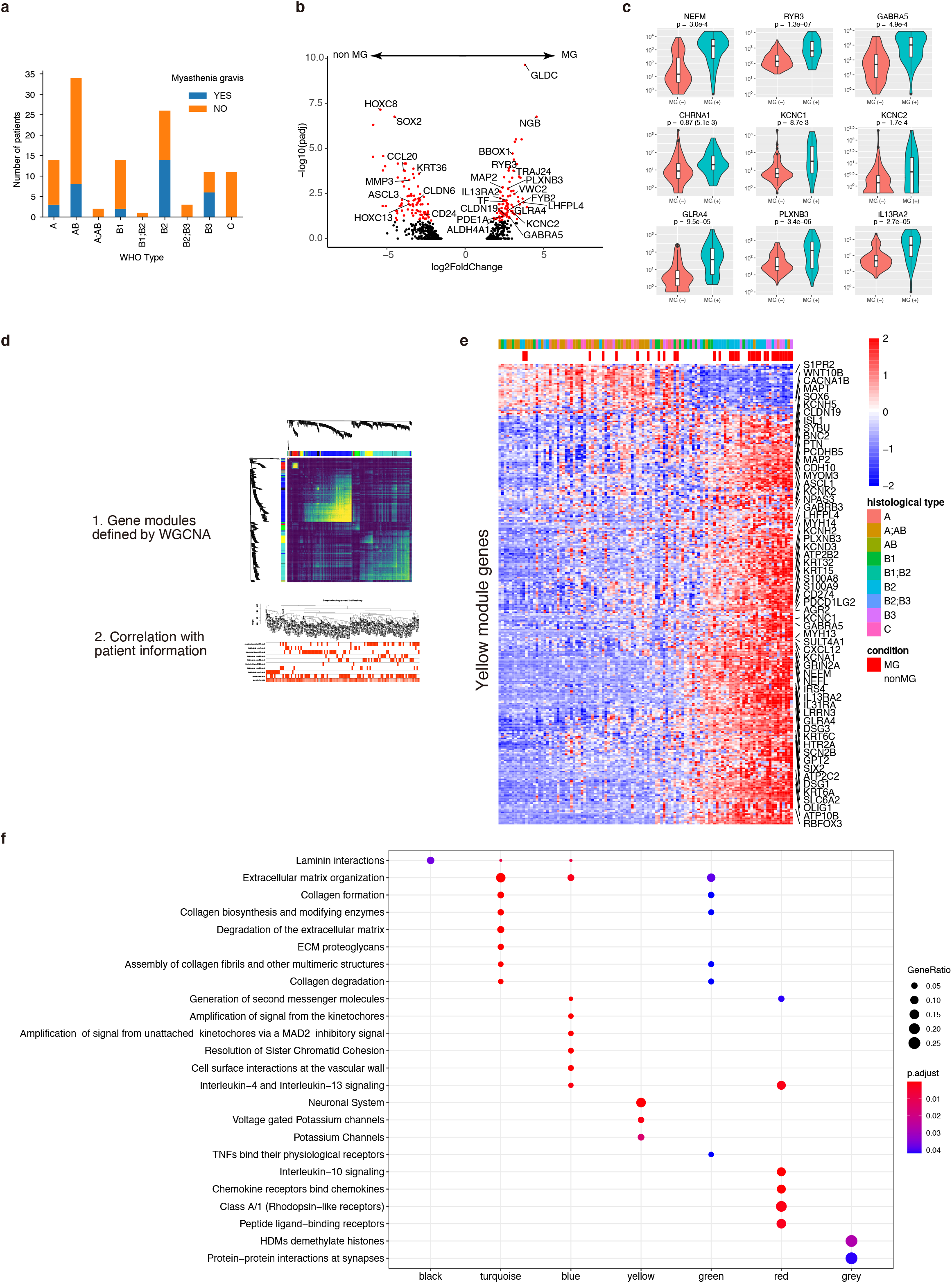
Global profiling of TCGA thymoma bulk RNA-seq dataset. a, Bar plot of patient distribution partitioned by WHO classification and MG status. b, A volcano plot showing adjusted *P*-value and *log*_*2*_ *fold change* for differential testing of genes between MG and non-MG patients in bulk RNA-seq of thymoma. Red dots represent statistically significant genes (*P*_*adj*_ < 0.1, |*log*_*2*_ *fold change*| > 1). c, Violin plots of DESeq2 normalized expression for MG-specific genes. Adjusted *P*-value by DESeq2 and for *CHRNA1 P*-value calculated by a two-sided Mann-Whitney *U* test (in parentheses) are shown. d, Workflow for Weighted Correlation Network Analysis (WGCNA). In WGCNA analysis, we first defined gene modules based on gene-wise expression correlation, then integrated clinical information. e, Heatmap showing standardized expression of genes in the yellow module. Samples were sorted by eigengene value. WHO classification and MG status are shown at the top of the heatmap. f, REACTOME pathways enriched in each module.

**Extended Data Fig. 3.**
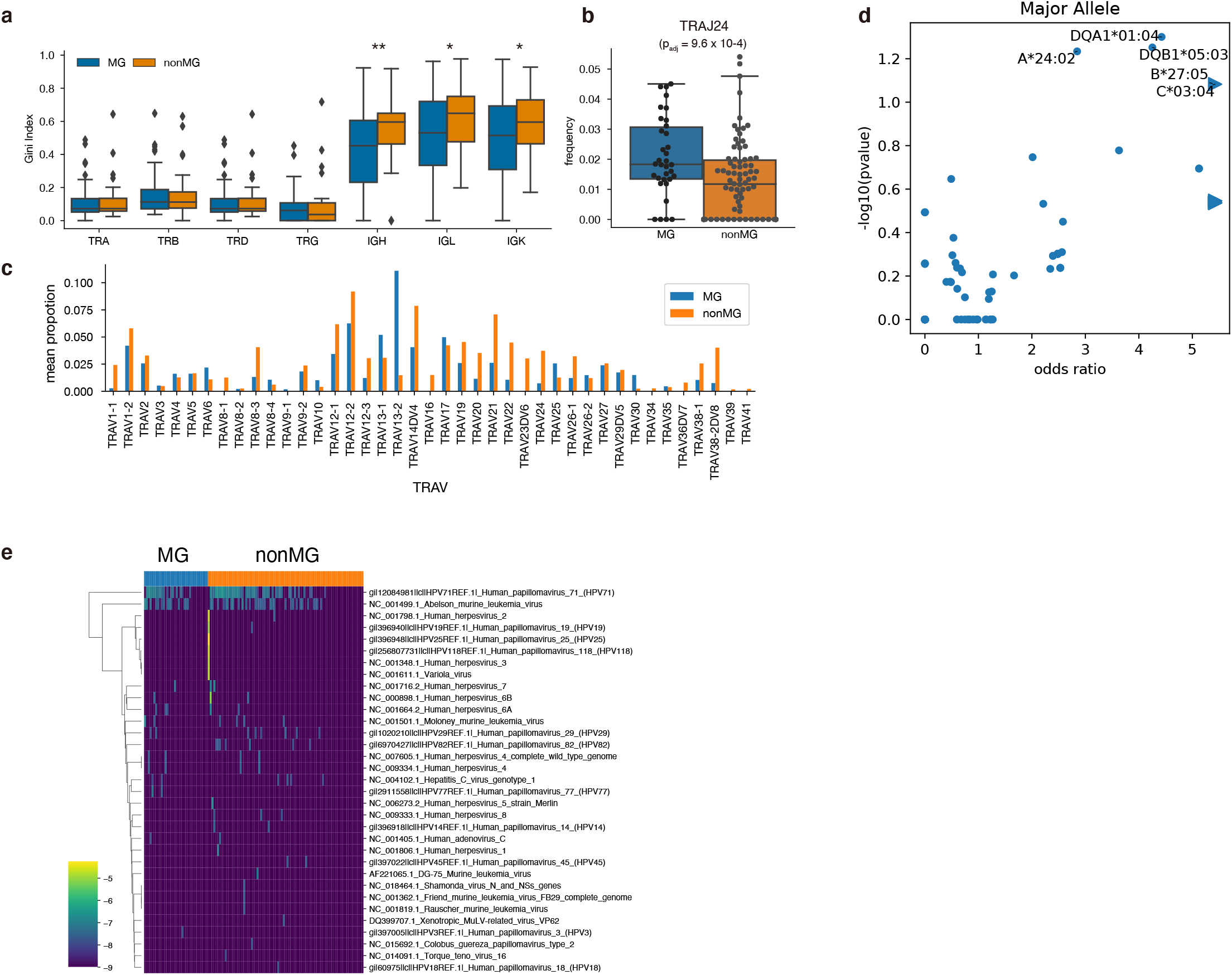
TCGA thymoma bulk RNA-seq dataset elucidated Immune characteristics of MG. a, Amino acid sequences diversity of immunoreceptors in CDR3 regions. The Gini index was used as an index of the complexity of repertoires in each group, and differences between MG and non-MG were tested using a two-sided Mann-Whitney *U* test. \**P* < 0.05, \*\**P* < 0.01. b, Box plot showing the frequency of *TRAJ24* in MG- and non-MG-thymoma. c, Bar plot of the frequency of TRAJ genes paired with *TRAJ24*. *TRAV13-2* was 7.50 times more frequent but not statistically significant. d, Volcano plot showing association of HLA major alleles with MG. Data were analyzed using a two-sided Fisher’s exact test. e, Heatmap of detected viruses in thymoma. The color indicates the number of transcripts mapped for each virus.

**Extended Data Fig. 4.**
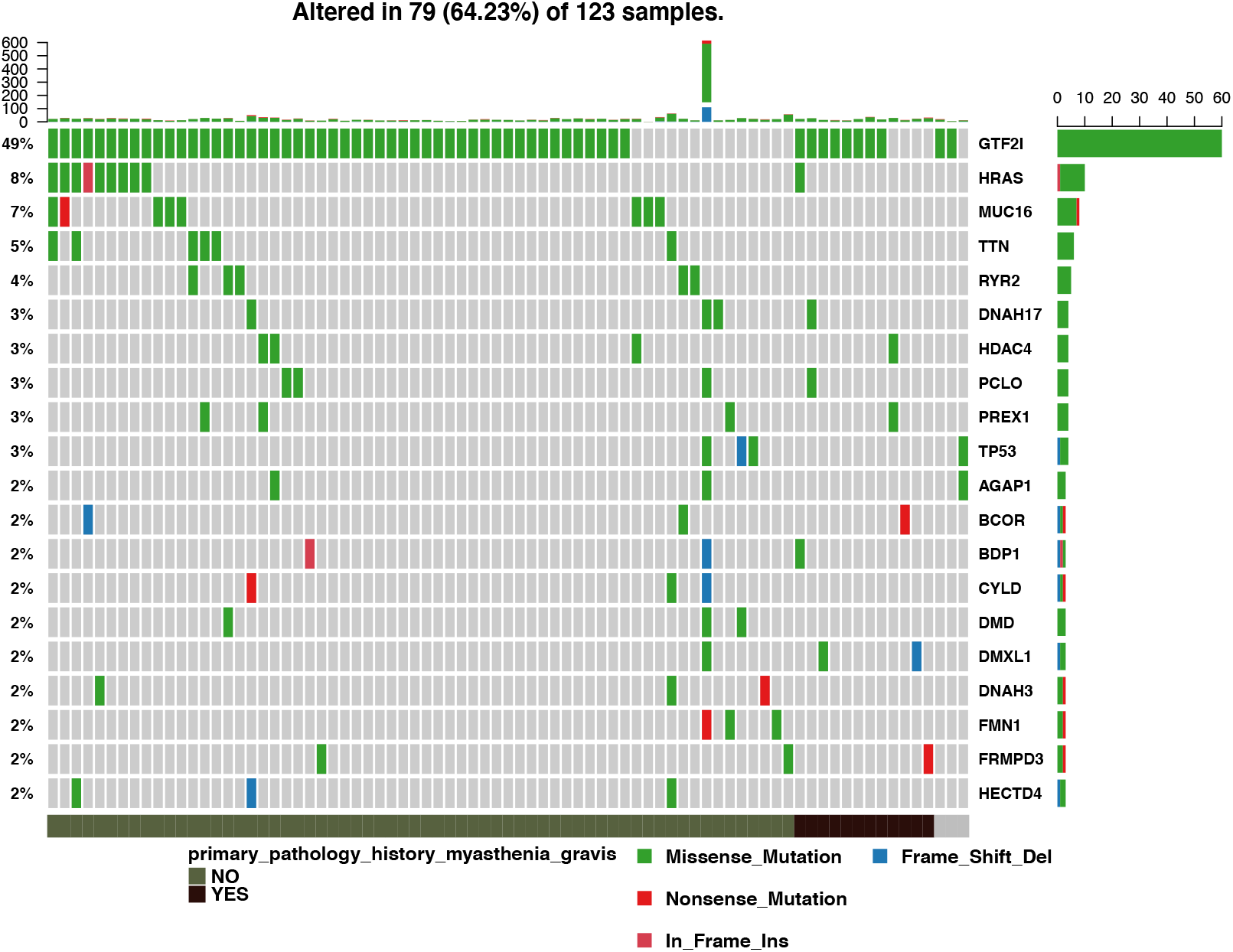
No somatic mutation in TCGA thymoma was associated with myasthenia gravis. Somatic mutations detected within TCGA thymoma samples with and without MG. Rows show observed variants aggregated by genes, and columns show individuals with MG status (below). The color represents the type of mutation. The frequency of the gene was mutated (right), and the abundance of mutation in each individual (top) is also shown.

**Extended Data Fig. 5.**
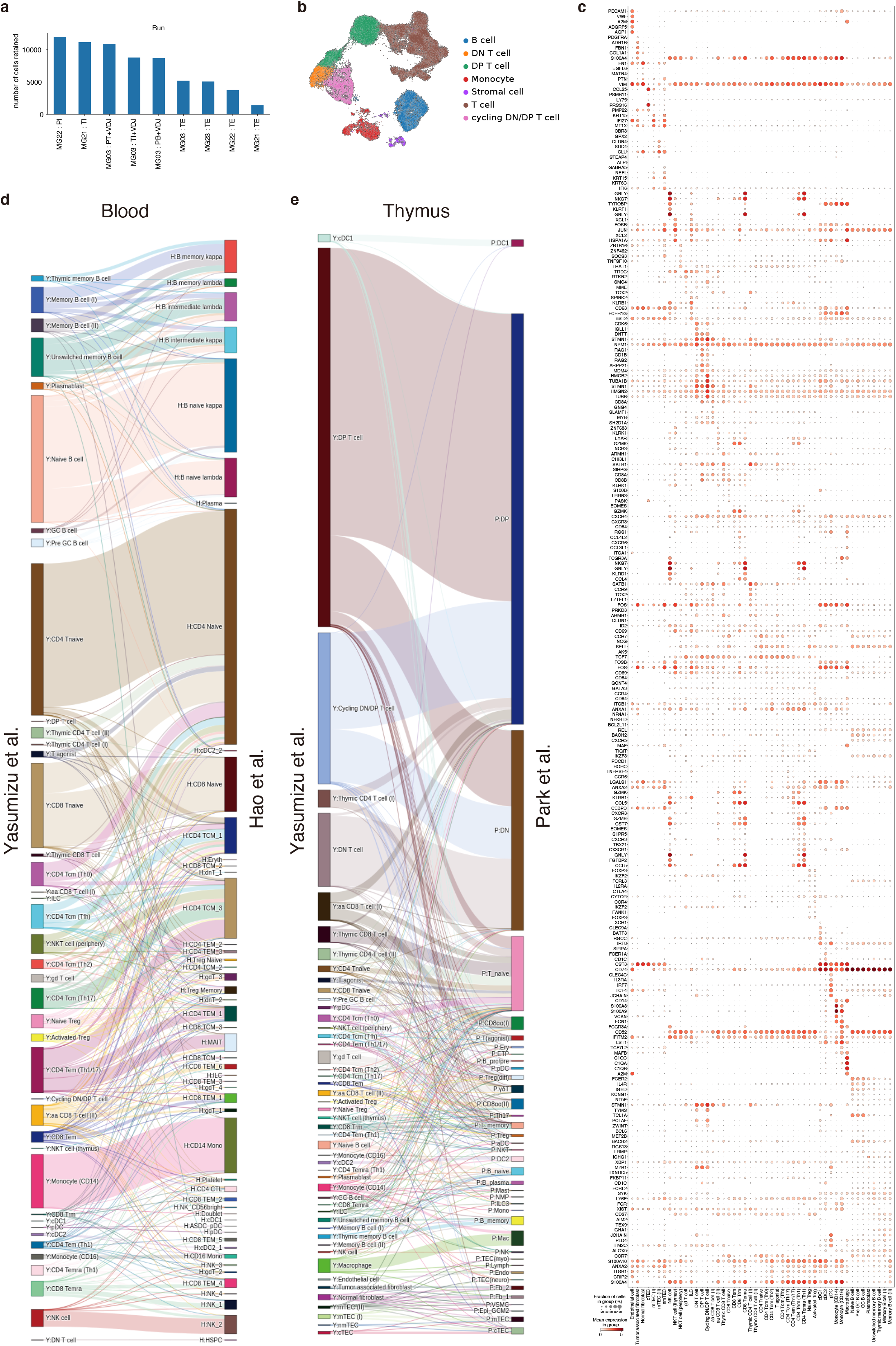
Confirmation of scRNAseq annotation and embedding. a, The number of cells recovered from each sample. We collected cells from four individuals with MG and thymoma. The abbreviation after the underscore in the sample ID indicates the source of the sample. TI: Thymoma immune cells, TE: Thymoma Epithelial cells, PI: Periphery immune cells, PT: Periphery CD4^+^ T cells, PB: Periphery CD19^+^ B cells. VDJ indicates 10x Genomics 5’+VDJ kit; otherwise, 10x Genomics 3’ GEM v3. b, The major categories on UMAP embedding. c, Detailed dot plot depicting signature genes’ mean expression levels and percentage of cells expressing them across clusters. d,e, Sankey diagrams showing cells aligned to each other in the thymus (d) and blood (e) of healthy individuals and our data set.

**Extended Data Fig. 6.**
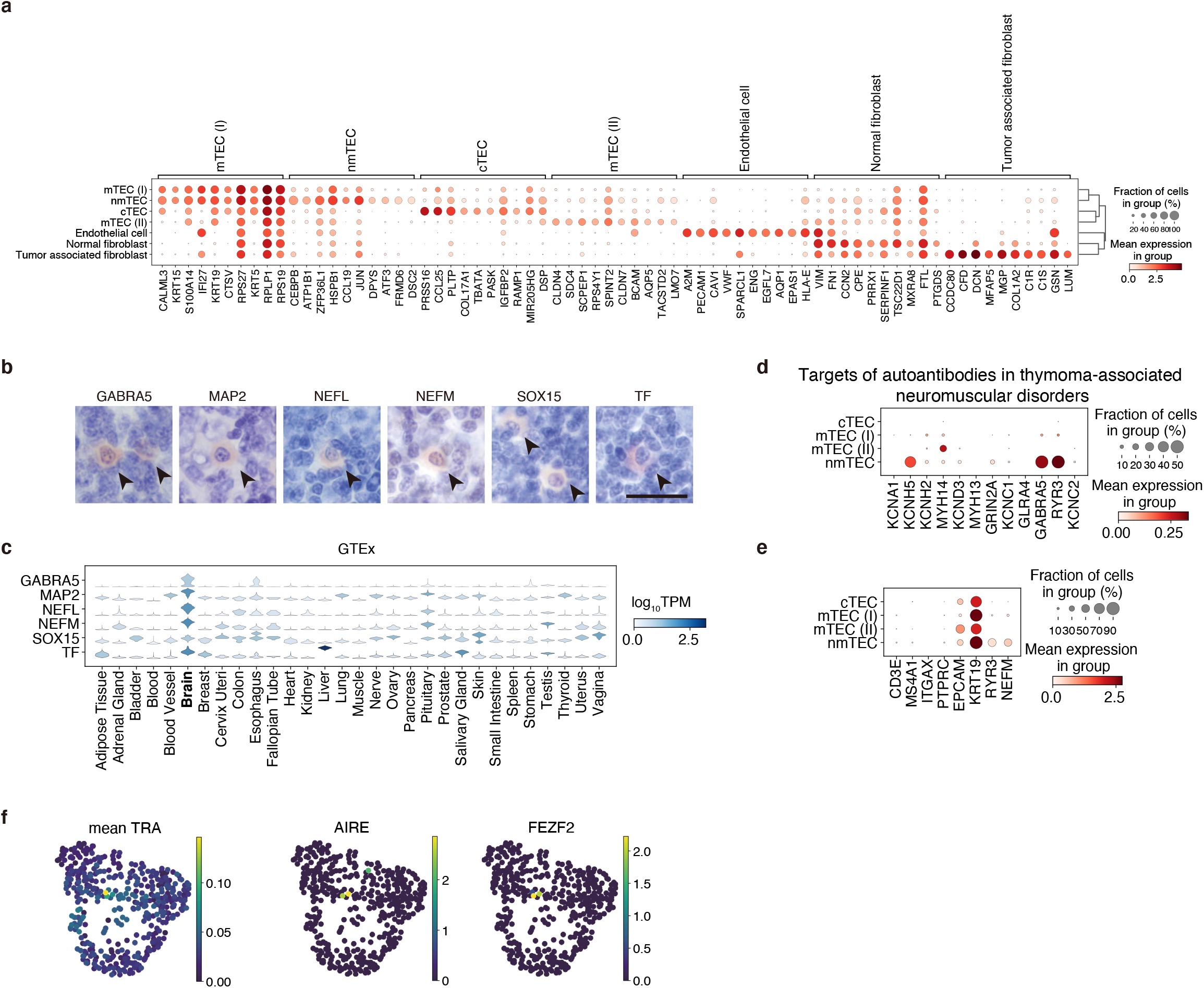
Detailed single-cell profiling of TEC cells. a, Dot plot of signature gene expression in stromal cell clusters. b, Immunohistochemistry of the yellow module genes. Scale bar: 20μm. c, Violine plots of the yellow module genes’ expressions across tissues in GTEx samples. d,e, Dot plot of gene expression of targets of autoantibodies in thymoma-associated neuromuscular disorders (d) and marker genes (e) in TEC clusters. f, UMAP embedding of mean expression of tissue-restricted antigens (TRAs) defined using GTEx bulk RNA-seq from systemic organs (left), and essential genes for TRA regulation; *AIRE* (middle), *FEZF2* (right).

**Extended Data Fig. 7.**
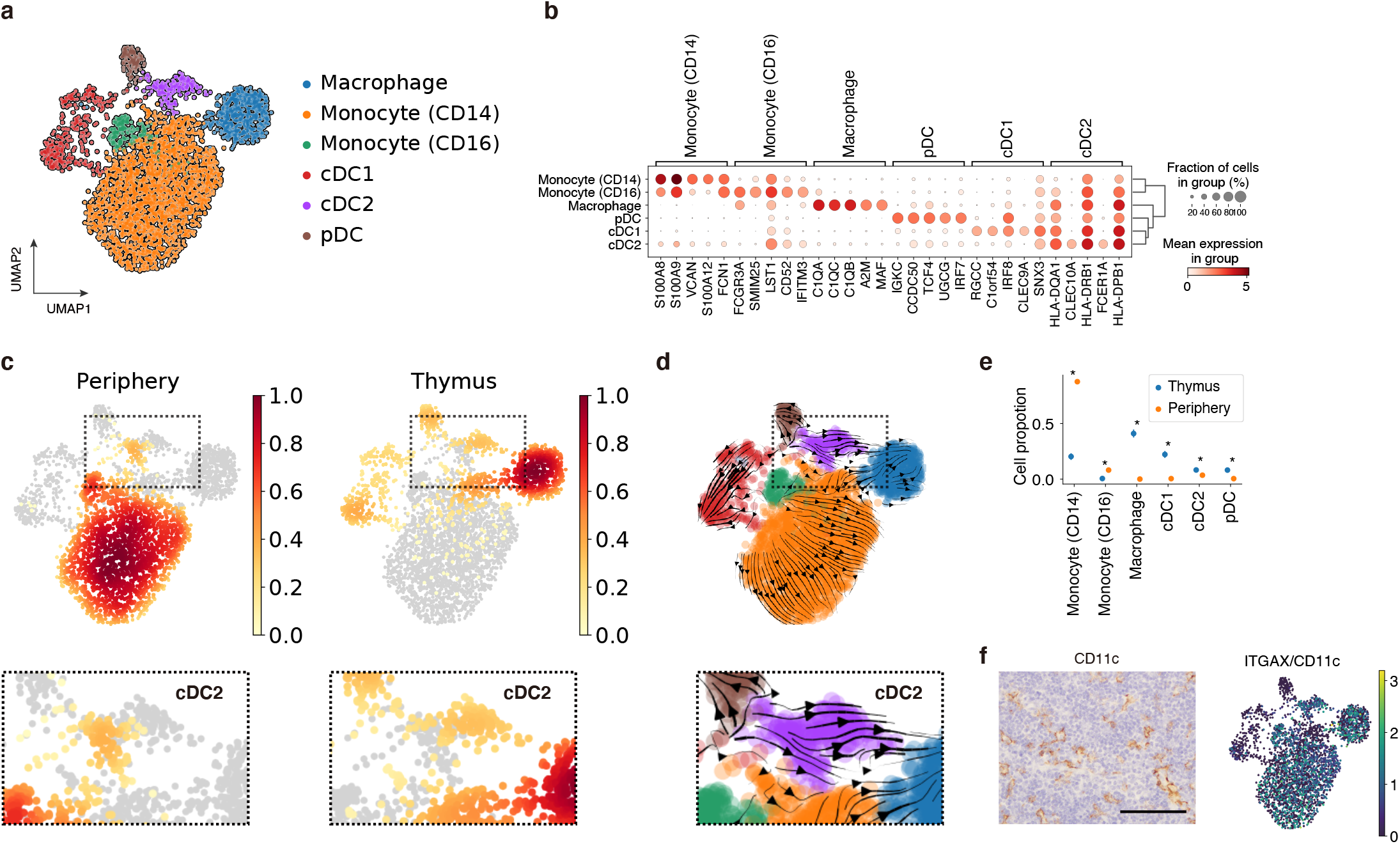
single-cell profiling of myeloid cells. a, UMAP embedding for myeloid cell clusters of thymoma and peripheral blood. b, Dot plot of gene expression of marker genes of each myeloid cell cluster. c,d, Density plots showing myeloid cell accumulation in the periphery (left) and thymus (right) (c) and RNA velocity in myeloid cells (d). Upper figures show the global picture, and lower images show the local picture focusing on cDC2s. e, Cell proportion of each myeloid cell cluster in thymoma and peripheral blood. \**FDR* < 0.05. f, Representative DAB staining for CD11c in MG-thymoma stained (left) and UMAP embedding of *ITGAX* (CD11c) expression. Scale bar: 100μm.

**Extended Data Fig. 8.**
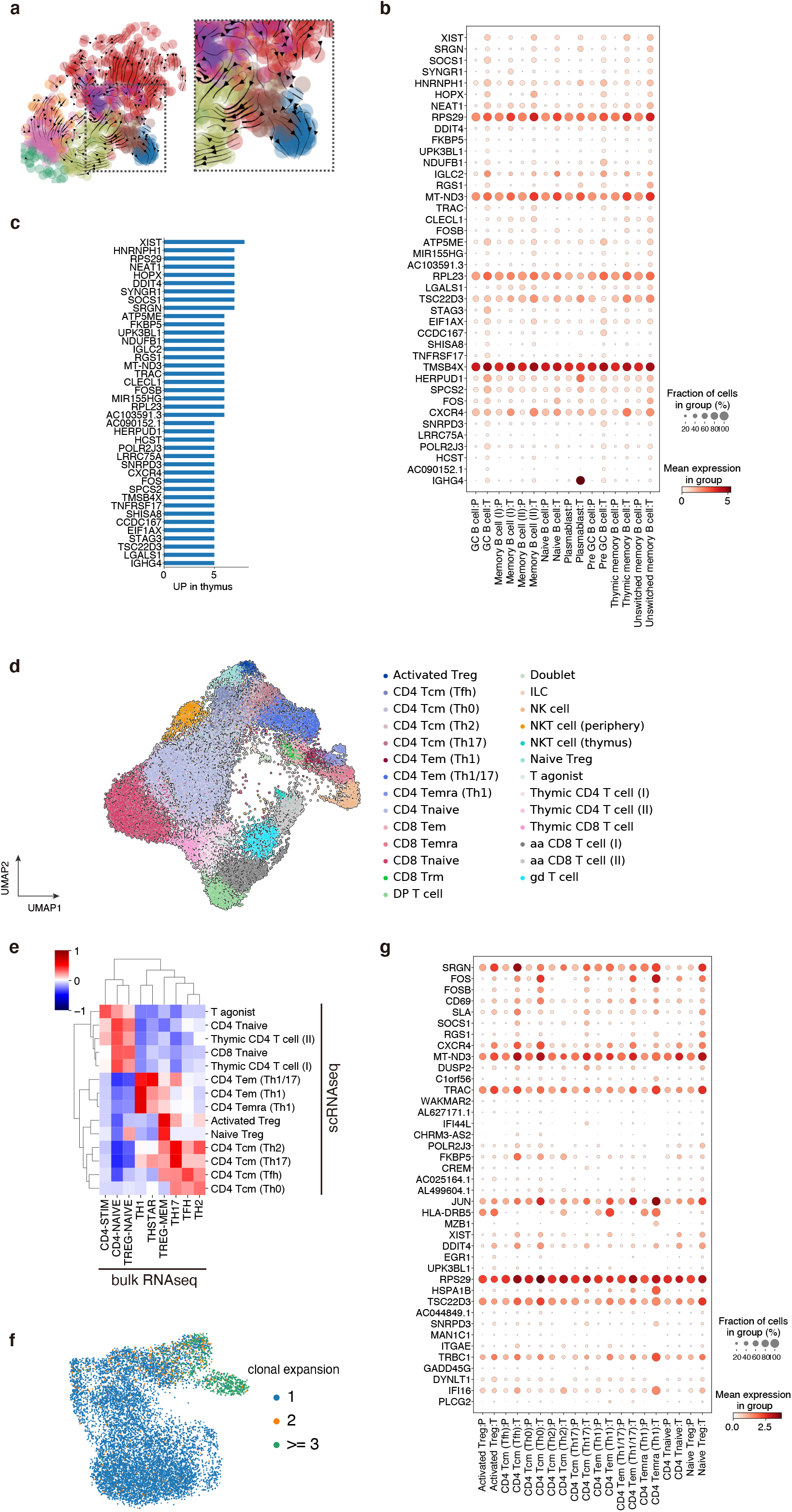
Detailed single-cell profiling of B cells and T cells. a, RNA velocity of intrathymic B cell in the global picture and local picture focusing on GC B cells and the neighboring cells. b, Dot plot of B cell gene expression of genes preferentially expressed in thymus across cell type. In column labels, P: peripheral blood, T: thymoma. c, Bar plot of thymus specific genes across B cell clusters ranked by the number of cell types where each gene was upregulated (*P*_*adj*_ < 0.05 and *log*_*2*_ *fold change* > 1) in B cells. d, UMAP embedding for T cell clusters except for DN and cycling DN/DP T cells of thymoma and peripheral blood. e, Heatmap of the correlation between CD4^+^ T cell clusters defined by our scRNAseq dataset and bulk RNA-seq sorted from peripheral blood established by the DICE (Database of Immune Cell Expression, Expression quantitative trait loci (eQTLs) and Epigenomics) project. f, UMAP embedding depicting the size of clonotypes. g, Dot plot of CD4^+^ T cell gene expression of genes preferentially expressed in thymus across cell type. In column labels, P: peripheral blood, T: thymoma.

**Extended Data Fig. 9.**
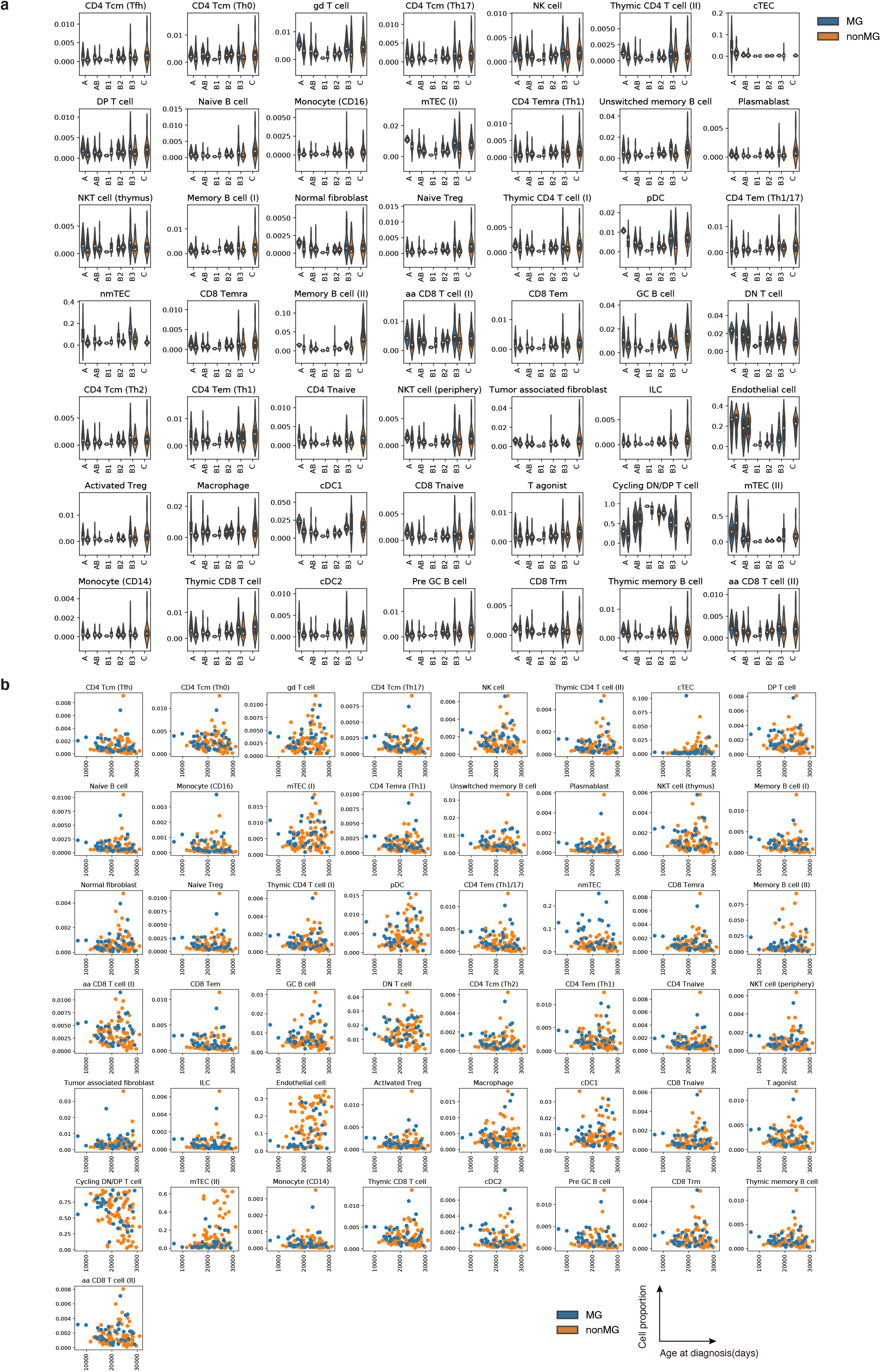
Inferred cell proportion in TCGA bulk RNA-seq in thymoma. a, Violin plots of deconvoluted cell proportion partitioned by WHO classification and MG status. b, Scatter plot showing the relationship between deconvoluted cell proportion (y-axis) and age at diagnosis (x-axis). Dot color represents MG status.

**Extended Data Fig. 10.**
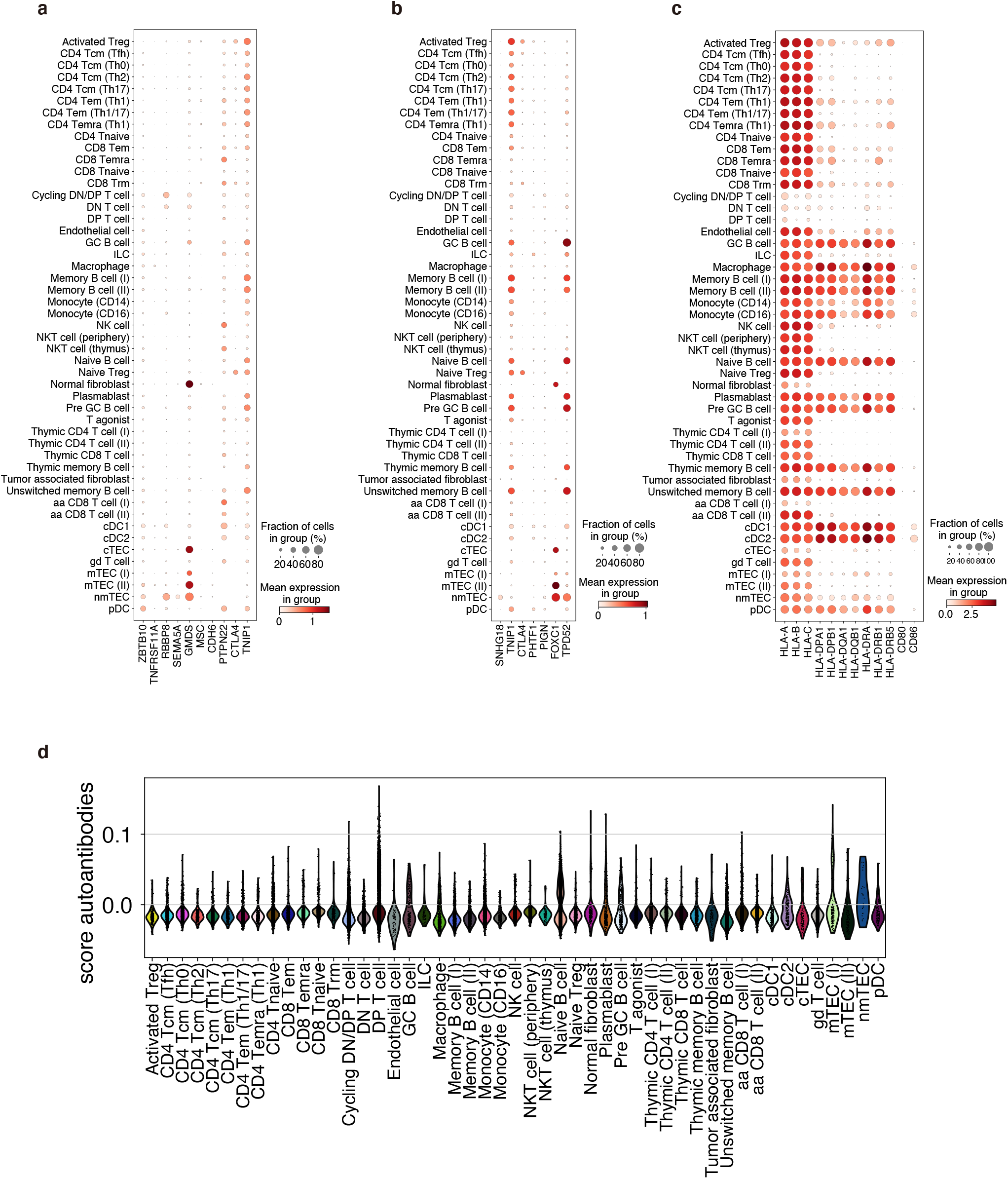
Cell-type wide expression of GWAS reported genes, HLA, and targets of autoantibodies thymoma-associated neuromuscular disorders. a-c, Dot plot of gene expression of GWAS reported genes (a), GWAS genes mapped by eQTL (b), and HLA and costimulatory molecules (c) across cell types. d, Violin plots of the signature score of targets of autoantibodies causing neuromuscular disorders associated with thymoma listed in Supplementary Table 6.

