## Supplementary Figures for "Myasthenia gravis-specific aberrant neuromuscular gene expression by medullary thymic epithelial cells in thymoma"

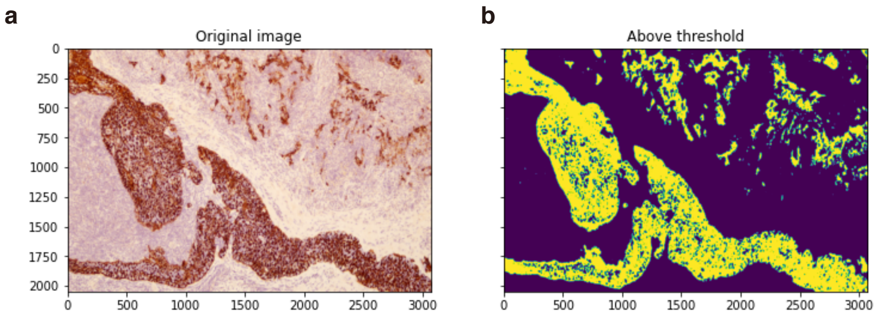


**Supplementary Figure 1** DAB signal segmentation and quantification. (a) An original DAB image under x10 objective. (b) The DAB positive area was determined automatically. We split color into three channels; DAB, hematoxyline, and eosin, then discriminated DAB positive area with an empirical threshold.

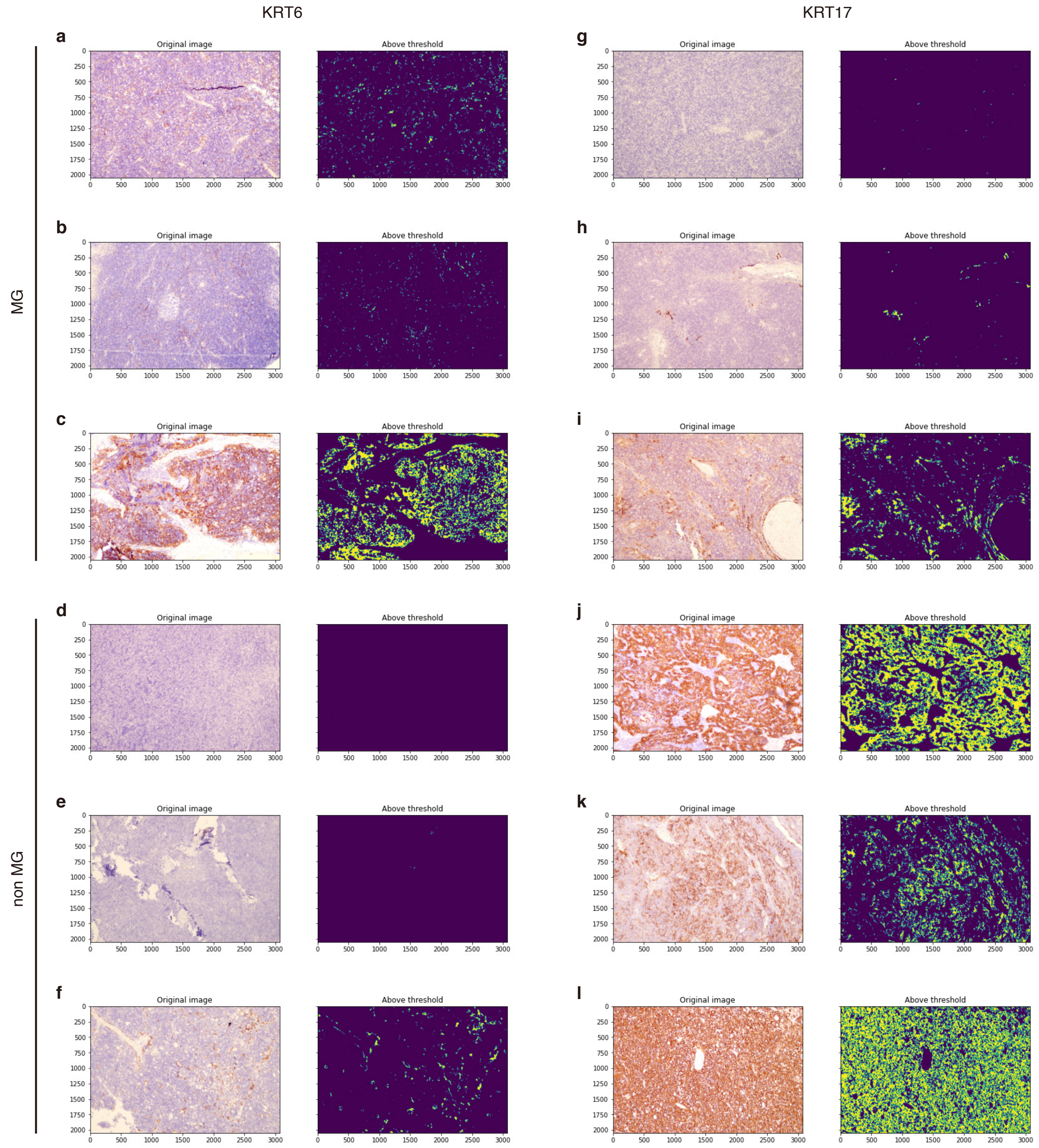


**Supplementary Figure 2** The representative images of KRT6 (**a**,**b**,**c**,**d**,**e**,**f**) and KRT17 (**g**,**h**,**i**,**j**,**k**,**l**) staining in MG (**a**,**b**,**c**,**g**,**h**,**i**) and non-MG (**d**,**e**,**f**,**j**,**k**,**l**) thymoma under x10 objective. **a** and **g**; **b** and **h**; **c** and **i**; **d** and **j**; **e** and **k**; **f** and **l** are tissue sections from corresponding donors.

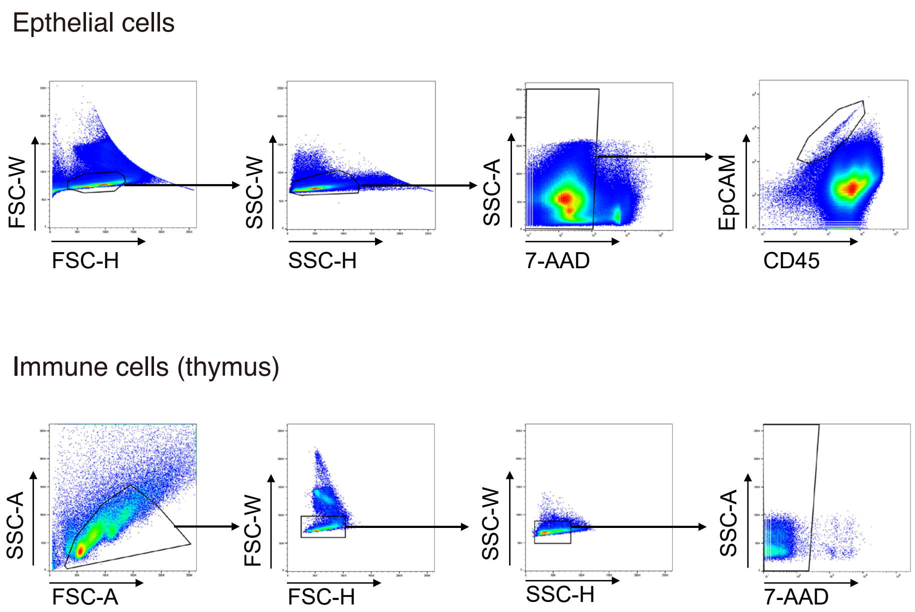


**Supplementary Figure 3** FACS gating for single-cell experiments.

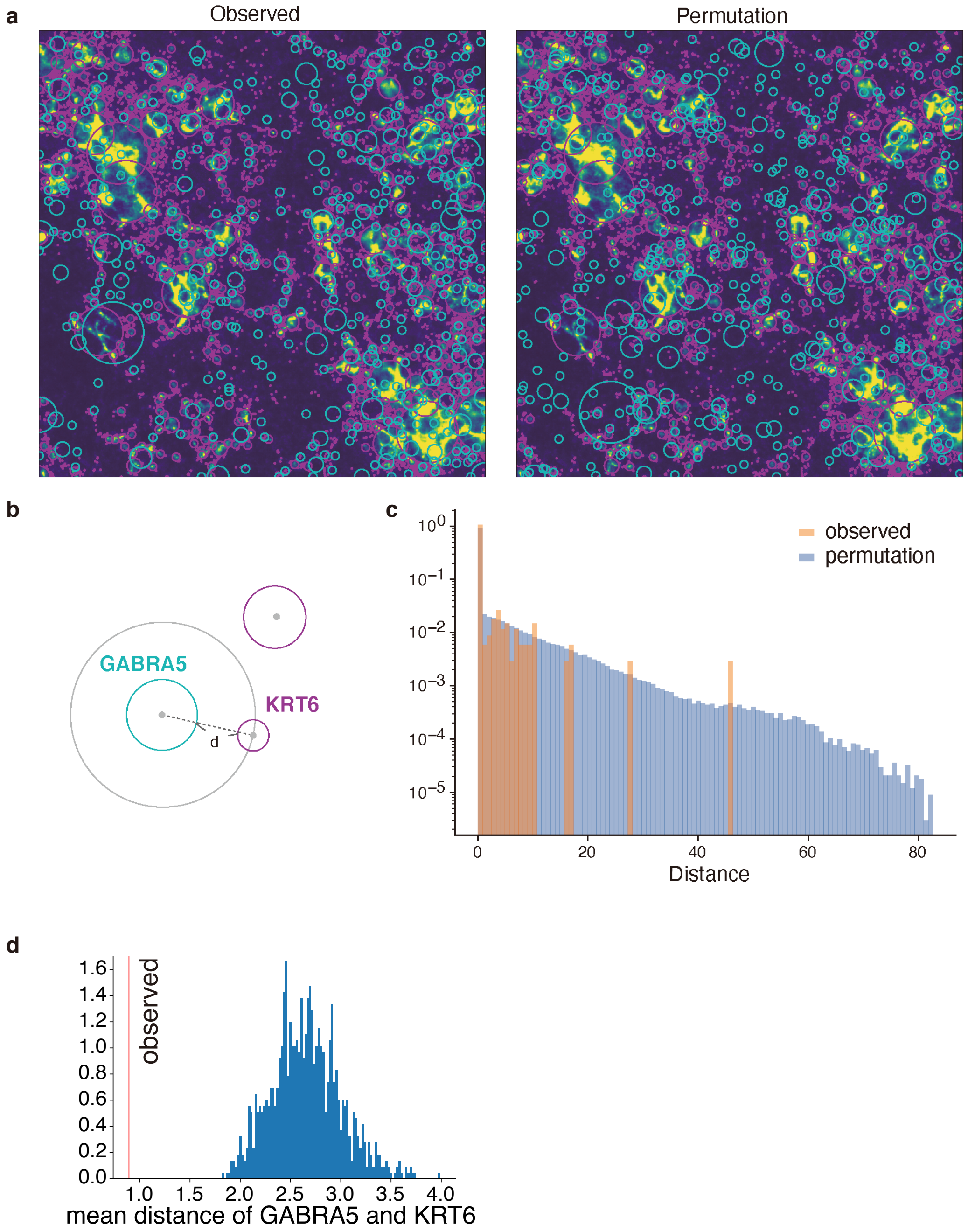


**Supplementary Figure 4** GABRA5 and KRT6 were co-localized. (a) Signal blobs and permutated blobs for GABRA5 (blue) and KRT6(purple). We conducted blob detection using immunofluorescence images stained GABRA5 and KRT6 (left). We also permutated blobs whose sizes and numbers were the same as the original image for 1000 sets (right). (b) The definition of distances of blobs. We measured the distance from GABRA5 blobs to the nearest KRT6 blobs. (c) The histogram is representing distances for observed blobs (orange) and permutated blobs (blue). (d) Mean distance of GABRA5 and KRT6 (a red line) and these for 1000 sets of permutation. The observed mean distance was smaller than all of the permutation sets.

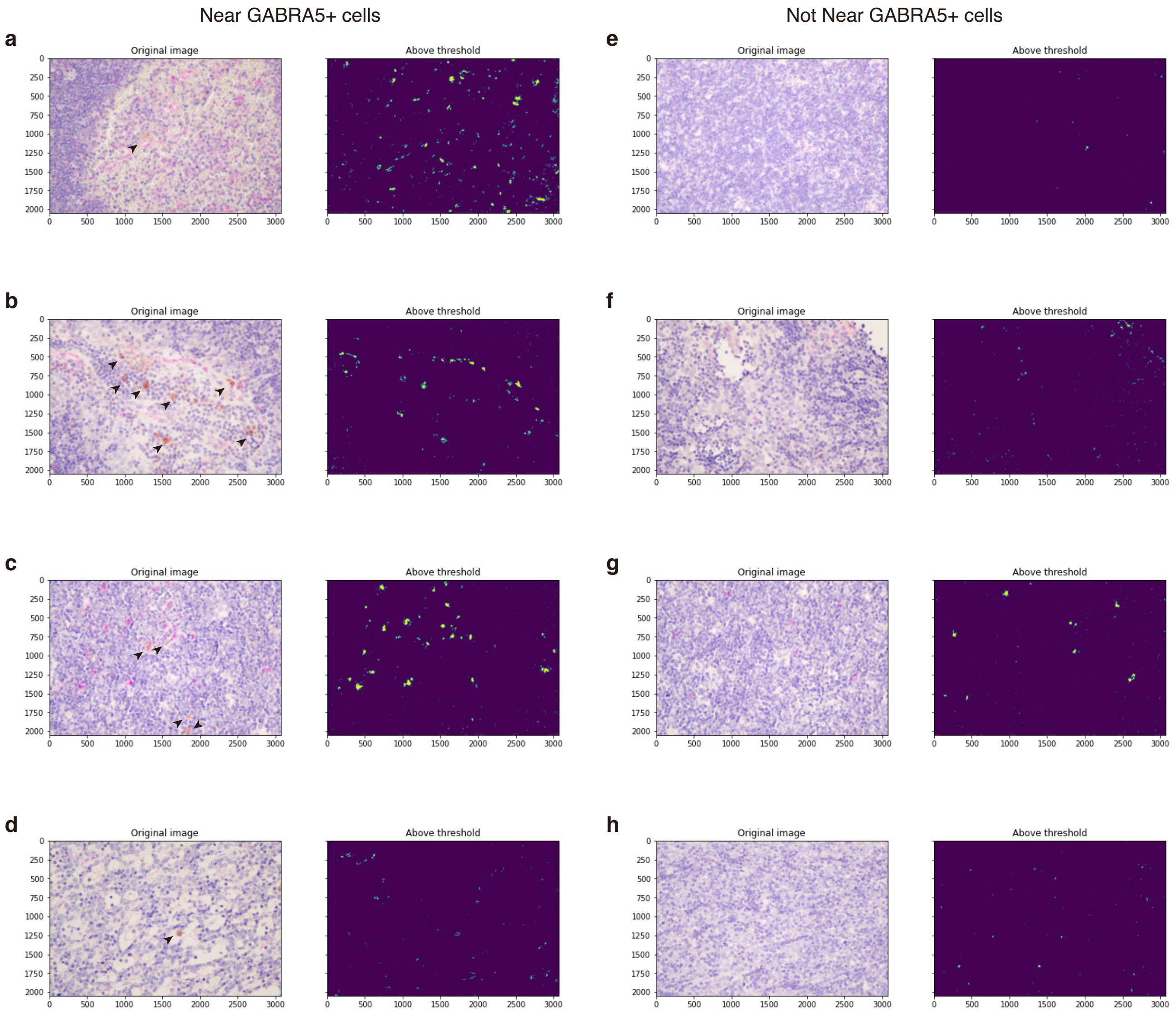


**Supplementary Figure 5** The representative images of CD31+ (purple) endothelial cells (left) and detected CD31 signals (right) near (**a**-**d**) and not near (**e**-**h**) GABRA5+ (DAB) cells under x40 objective. **a** and **e**; **b** and **f**; **c** and **g**; **d** and **h** are tissue sections from corresponding donors. Arrowheads represent GABRA5+ cells.

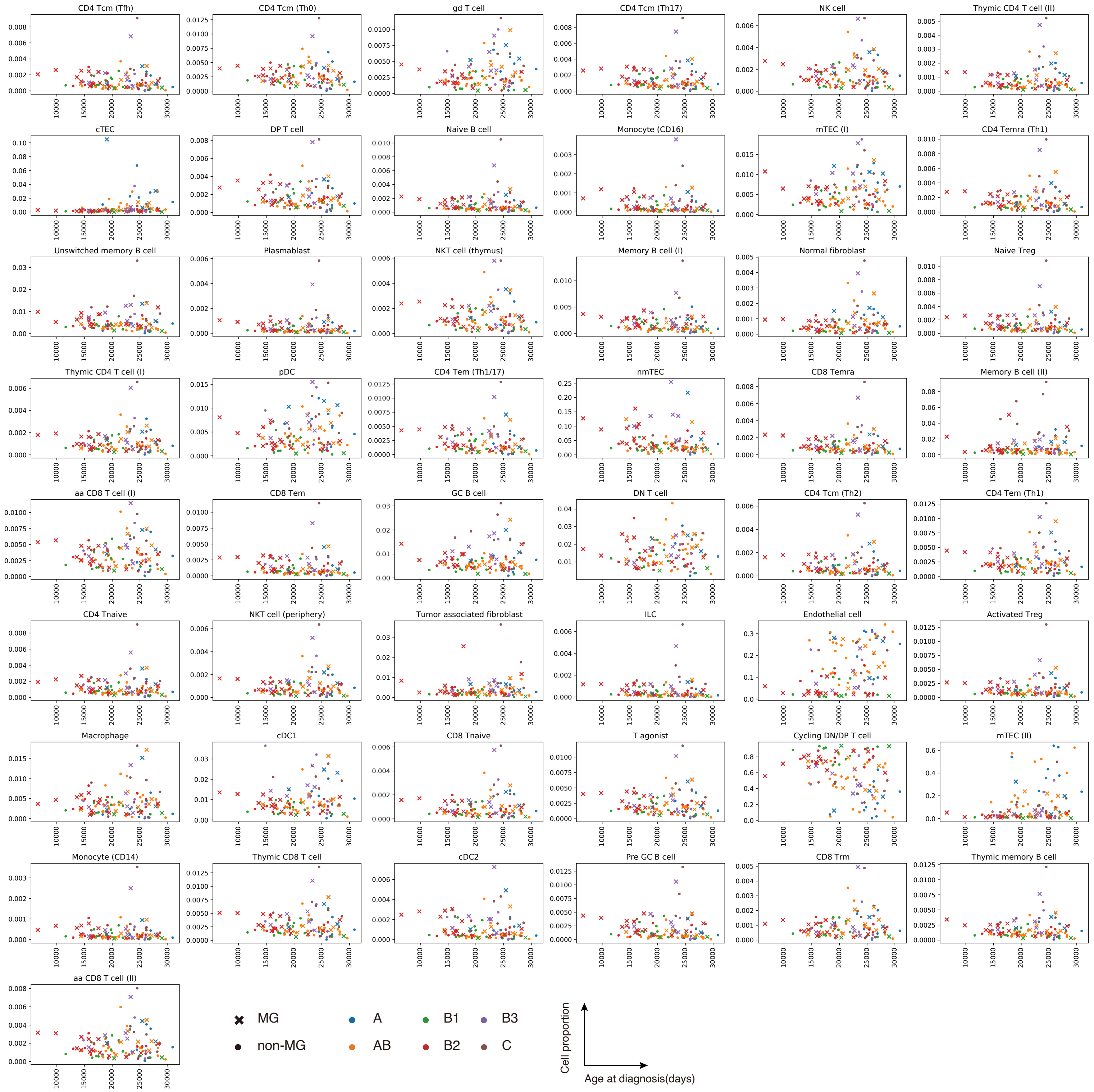


**Supplementary Figure 6** The scatter plot showing deconvoluted cell proportion and age at diagnosis. The marker shape represents the disease status, and the color does WHO classification.
